# Single cell profiling of CRISPR/Cas9-induced OTX2 deficient retinas reveals fate switch from restricted progenitors

**DOI:** 10.1101/538710

**Authors:** Miruna G. Ghinia Tegla, Diego F. Buenaventura, Diana Y. Kim, Cassandra Thakurdin, Kevin C. Gonzalez, Mark M. Emerson

## Abstract

Development of the vertebrate eye, like many developmental systems, depends on genes that are used iteratively in multiple distinct processes. The OTX2 transcription factor is one such gene, with a requirement for eye formation, photoreceptor formation, and retinal pigment epithelium specification, among others. Recent evidence has suggested that OTX2 is also expressed in subsets of retinal progenitor cells with restricted fate choices. However, given the multiple roles for OTX2 and limitations of conventional conditional knockout strategies, the functional significance of this expression is unknown. Here we use CRISPR/Cas9 gene editing to produce mutations of OTX2, identifying similar phenotypes to those observed in human patients. In addition, we use single cell RNA sequencing to determine the functional consequences of OTX2 gene editing by CRISPR/Cas9 on the population of cells derived from OTX2-expressing retinal progenitor cells. We not only confirm that OTX2 is required for the generation of photoreceptors, but also for maintaining the proliferative potential of cells and suppressing the formation of specific retinal fates. These include subtypes of retinal ganglion and horizontal cells normally associated with these progenitor types, suggesting that in this context OTX2 functions to repress sister cell fate choices. Upregulation of key transcription factors involved in the formation of these cells was observed suggesting that OTX2 is upstream of critical nodes of gene regulatory networks of these alternative fates.

## Introduction

The vertebrate retina is a highly structured tissue with five neuronal cell classes, each comprising multiple cell types, and one type of glia. The specification of these diverse types is a dynamic process that is conserved across species, with the commitment to a final fate of retinal ganglion cells (RGCs), cone photoreceptors (PRs) and horizontal cells (HCs) in the first developmental wave, followed progressively by amacrine cells (ACs), rod PRs, bipolar cells (BCs) and, lastly, Müller glia (Masland 2001, Bassett and Wallace 2012).

Elucidation of the mechanisms by which each cell type is generated from multipotent retinal progenitor cells (RPCs) is crucial for the development of successful cell replacement therapies for patients that suffer from retinal degeneration. Initial studies using viral labeling showed that multipotent RPCs have the potential to give rise to all retinal cell types, in a mechanism that can be explained by both deterministic and probabilistic models (Turner and Cepko 1987; Cepko et al., 1996; Vitorino et al., 2009; He et al., 2012). Recently, identification of markers of restricted lineages enabled the characterization of these types of RPCs. For example, one class of restricted RPCs gives rise to all cell types except RGCs (Brzezinski et al., 2011), another produces all cell types except RGCs and Müller glia (Hafler et al., 2012), and a lineage that is largely restricted to only two cell types – cone PRs and HCs (Emerson et al., 2013).

This last lineage is defined by the activity of the cis regulatory module 1 (CRM1) of the gene THRB. Previous studies have determined that the activation of the THRBCRM1 element requires the expression and direct binding of the OTX2 (orthodenticle homeobox 2) and ONECUT1 transcription factors (Emerson et al., 2013). Moreover, a recent study has identified conserved cellular and molecular changes that occur during the generation of these fate-restricted RPCs, but not in multipotent RPCs (Buenaventura et al., 2018). However, the roles of OTX2 and ONECUT1 in the establishment of this restricted signature are not well understood.

OTX2 is one of the key regulators of nervous system development, as it is involved in forebrain and midbrain specification (Acampora et al., 1995; Ang et al.,1996, Simeone et al., 2002), pineal and pituitary gland development (Nishida et al., 2003; Henderson et al., 2009), as well as development of sensory structures such as the inner ear (Matsuo et al., 1995) and visual system (Beby et al., 2013). In the visual system, OTX2 is expressed in different subsets of cells as development progresses –first, during optic vesicle formation, OTX2 directs evagination of the optic vesicle to contact the surface ectoderm along with RAX, PAX6, HES1 and SIX3 (Adler and Canto-Soler 2007). Later, during the specification of the neuronal and retinal pigment epithelium (RPE) territories, OTX2 is highly expressed in the RPE (Martinez-Morales et al., 2001) and in a subset of RPCs that generate primarily cones and HCs (Emerson and Cepko, 2011, Emerson et al., 2013, Buenaventura et al., 2018), then in postmitotic PRs during the specification of retinal neuronal cell types (Nishida et al., 2003), in the PRs and BCs in the mature retina (Koike et al., 2007; Kim et al., 2008), and in a subset of Müller glia (Brzezinski et al., 2010). As a consequence of its involvement during early eye genesis, several studies have reported OTX2 mutations in humans with ocular malformations. The clinical manifestations range from unilateral and bilateral anophthalmia, microphthalmia, optic nerve aplasia to various forms of coloboma (reviewed by Gat-Yablonski 2015).

In addition, previous studies have examined the role of OTX2 in the development of the neuronal retina using conditional floxed mouse models, as homozygous OTX2 mutants die embryonically due to defects in the specification of the anterior neuroectoderm (Acampora et al., 1995). When OTX2 loss is mediated by a post-mitotic PR Cre driver, a severe loss of PRs was observed, while the number of PAX6-positive cells was increased, suggesting a trans-differentiation of the mutant cells into amacrine-like neurons (Nishida et al., 2003). Microarray analysis of these retinas has supported this conclusion (Omori et al., 2011). Similar PR loss and abnormal generation of amacrine-like cells, along with loss of BCs and HCs was reported when OTX2 ablation was initiated throughout the early retina (Sato et al., 2007). Despite the contributions from these studies in implicating OTX2 as a crucial regulator of PR development, several questions remain unanswered. Specifically, what are the cell fates and underlying gene regulatory networks that are impacted by loss of OTX2? While previous analyses have suggested that ACs are ectopically induced upon loss of OTX2, these effects were characterized in the postnatal retina leaving the primary effects of OTX2 loss unknown. In addition, it has been determined that OTX2 expression is initiated in specific RPC populations prior to the formation of PRs and this expression would not be expected to be compromised using the previously used PR Cre drivers (Emerson et al., 2013; Buenaventura et al., 2018). Thus, it is unclear what the function of OTX2 is in these RPCs that will predominantly generate photoreceptors.

Here we report a characterization of OTX2 mutants during various stages of retinal development with a focus on OTX2-positive RPCs. CRISPR-induced OTX2 loss-of-function mutations were introduced in the context of the developing chicken retina *in vivo* and *ex vivo*, providing a robust and facile experimental system to determine OTX2 function. Successful ablation of OTX2 was demonstrated by immunofluorescence, gene expression and deep sequencing of the OTX2 gene. The effects of OTX2 loss on OTX2-positive RPCs were analyzed using a reporter active in this cell population combined with the examination of known markers and characterization using single cell RNA sequencing. We confirmed the strict requirement of OTX2 for the specification of PR cells. In addition, we determined that there was an induction of the formation of specific subtypes of RGCs and HCs from the OTX2 mutant population, suggesting that alternative fates are not randomly chosen, but are selected from those normally associated with OTX2-positive RPCs. This high-resolution analysis of OTX2 function provides new insights into the role of OTX2 PR formation in the vertebrate retina and the mechanisms of retinal cell fate choice.

## Results

### CRISPR/Cas9-induced ablation of the OTX2 gene yields severe anatomical defects

To generate mutations that disrupt the function of the OTX2 gene, three RNA guides were designed starting at mRNA positions 38, 140 and 166 in the coding sequence of the OTX2 gene, called guide 1 (g1), guide 2 (g2) and guide 3 (g3) for simplicity (Fig. 1A-B). All three guides were designed to target the homeodomain region or upstream of it. *In vivo* electroporation of OTX2 g2 at E1.5 (HH9-HH11) yielded severe anatomical defects of the electroporated eye upon analysis at E5, a timepoint when both the neuronal retina and the RPE territories have already been specified. For clarity, eyes electroporated with a plasmid carrying the sequence of the RNA g2 are herein called OTX2^CRISPR^ mutants.

**Figure 1.**
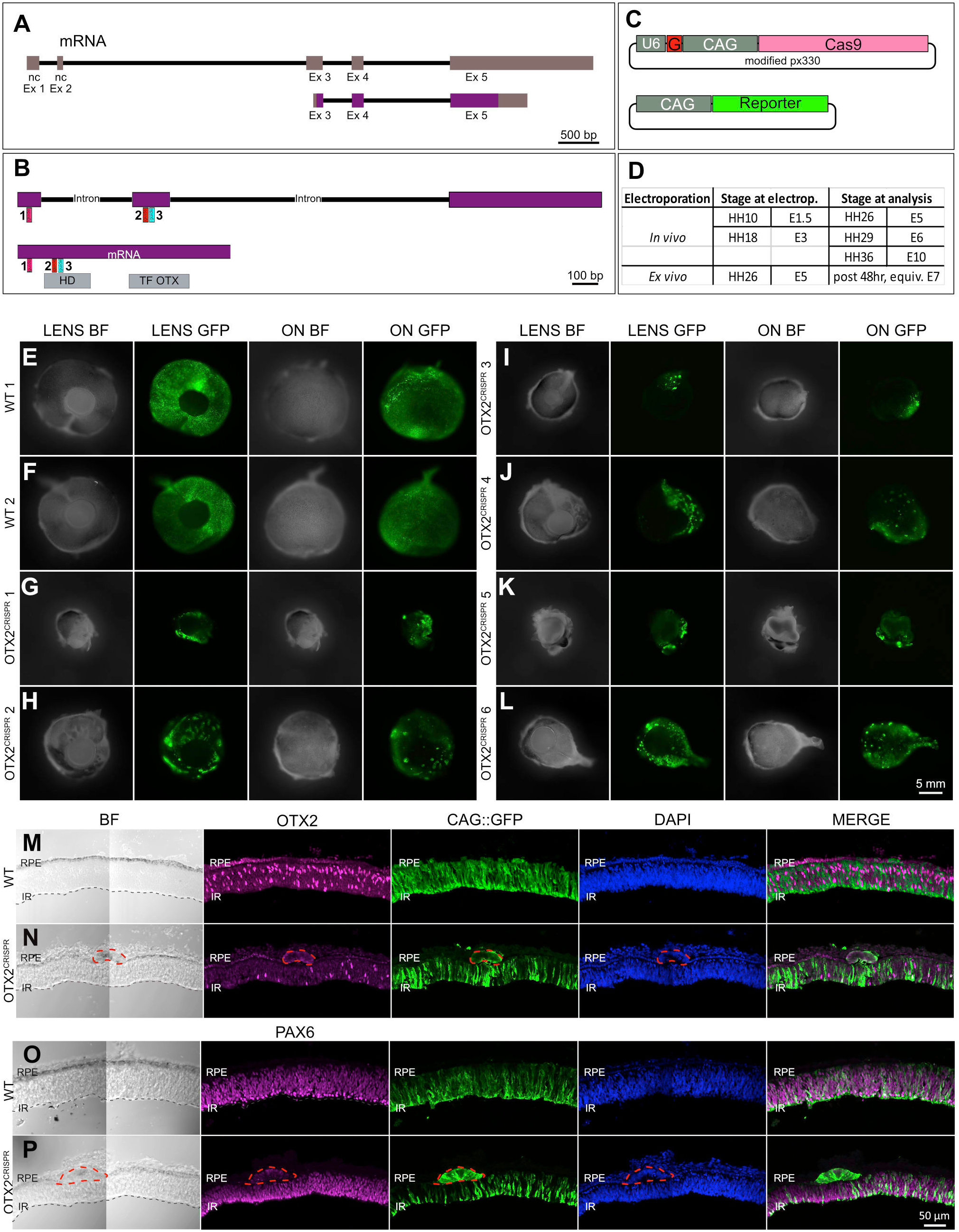
OTX2^CRISPR^ guide design and targeted electroporation of the chick eye cup yields severe abnormalities of the mutant eyes. **A**. Schematic representation of the OTX2 genomic locus in the *Gallus gallus* genome. Purple blocks represent the coding region. Gray blocks represent non-coding exon regions. **B**. Location of the guides 1-3 relative to the unspliced (top) and spliced (bottom) OTX2 mRNA. Grey boxes show the mRNA regions that encode the homeobox domain. **C**. Schematic of co-electroporated plasmids. U6 is the promoter for the guide RNA, denoted by G., CAG drives expression of Cas9 and fluorescent reporters. **D**. Time points for electroporation of CRISPR plasmids and analysis. **E-L**. Phenotypes observed after eye cups were electroporated with CRISPR Cas9 and guide 2 at E1.5/HH 10 and analyzed at E5/HH 26. Images were acquired from the frontal (LENS) and dorsal (ON) view of whole eyes. Green signal confirms electroporation efficiency of the CAG::GFP control plasmid. All mutants (**G-L**) display different degrees of microphthalmia and RPE depigmentation, while few show coloboma-like defects (**I, J, L**). **E** and **F** were electroporated with an empty p18 vector and the electroporation control plasmid and, are considered wild-type for OTX2. **M-P** Variation in retinal thickness was observed in mutants. OTX2 protein expression is severely diminished in mutants (**N)** as compared to WT (**M**). Mutant RPE is depigmented and abnormal cell clumps with strong GFP and no OTX2 are formed. RPE structures in mutants are PAX6 positive (dotted lines, **P**). Ex, Exon; nc, non-coding; HD, homeodomain; BF, brightfield; ON, optic nerve; RPE, retinal pigment epithelium; IR, inner retina.

The morphological defects observed in the OTX2^CRISPR^ mutants ranged from microphthalmia (Fig. 1G-L), patches of depigmentation in the RPE (Fig. 1H-J, L), defects of the optic nerve (ON), such as ON head enlargement and ON aplasia in few mutants, as well as both iridial and optic stalk coloboma-like phenotypes (Fig. 1L). For all embryos, the contralateral non-electroporated eye appeared to be normally developed (data not shown). The WT controls were electroporated with CAG::GFP as well as a p18 plasmid, which is a modified px330, that contained the Cas9 ORF driven by CAG but without a sgRNA sequence. In contrast to the OTX2^CRISPR^ eyes, the control eyes showed no morphological defects (Fig. 1E-F).

The mosaicism observed in the patches of depigmentation in the mutant RPE as well as the variable size of the mutant eyes reflects the electroporation efficiency as well as the fact that the mutations were induced through the non-homologous end-joining (NHEJ) after the Cas9 double-stranded break, and, therefore, the clones deriving from electroporated cells are different with respect to their targeted mutation.

### OTX2 mutations induced at E5 disrupt both the RPE and neuronal retina

To assess the structural defects observed in the OTX2^CRISPR^ mutants, a characterization using immunofluorescence was performed on vertical sections. Mutants varied in thickness of both RPE and neuronal retina (Fig. 1M-P and Suppl. Fig. 1). The RPE layer was found to contain tumor-like structures composed of cells that were depigmented, enlarged, intensely GFP-positive, and OTX2-negative. These structures were EdU negative and predominantly PH3-negative (Suppl. Fig. 1C,D,H,I), while ectopically expressing low levels of VSX2 (Suppl. Fig.1A,B), but high levels of PAX6 (Fig. 1P), suggesting a potential switch from RPE to neuronal fate.

At the developmental time of analysis, E5, OTX2-positive cells in the neuronal retina are normally observed in a dispersed pattern in the retina, with a population that strongly expresses OTX2 located in the outermost part of the retina and a more weakly expressing one, predominantly composed of RPCs, in the central retina (Buenaventura et al., 2018). The OTX2^CRISPR^ mutants displayed varying degrees of effects on OTX2 expression, ranging from absence of the outer OTX2 population to complete loss of OTX2 throughout the entire retina extent (Fig. 1M,N and Suppl. Fig. 1E,F,G).

### OTX2 mutations in the neuronal retina at different developmental timepoints

The mutations induced during optic cup formation at E1.5 yielded a broad range of morphological defects due to the absence of OTX2 in both neuronal retina and RPE, and therefore, made the function of OTX2 during PR development difficult to assess. To more specifically target the neuronal retina, *in vivo* electroporation experiments were performed at E3 (HH29), when the RPE and neuronal territories are already specified and the majority of cells are RPCs with high division potential, ensuring the propagation of the mutations to many progenies.

The CAG::GFP plasmid was used as electroporation control to trace the targeted cells. At this stage, the electroporated area is limited due to the nature of the experiment (Methods), and the sparse clones of mutated cells can therefore be analyzed in the context of a mostly unaffected retina. When mutant retinas are analyzed at E6, the total number of OTX2-positive cells doesn’t appear overtly affected, however, the GFP-positive cells generally lack OTX2 protein (Suppl. Fig. 2A,B).

**Figure 2.**
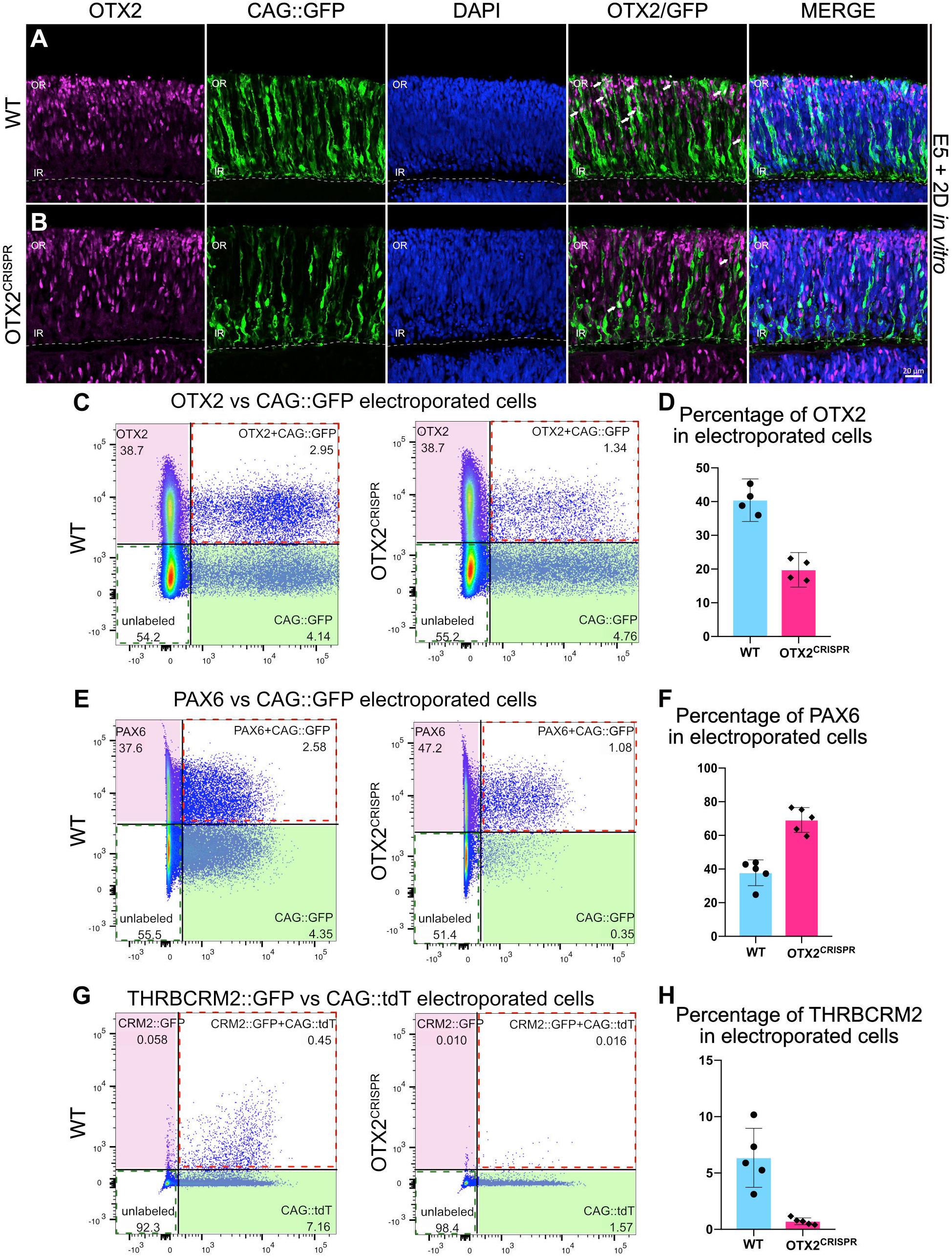
Assessment of OTX2^CRISPR^ retinas electroporated at E5 and analyzed post 48h. **A-B**. Confocal microscopy assessment of vertical sections of retinas electroporated with CAG::GFP and immunostained for OTX2 (magenta), GFP (green) and DAPI (blue) (**A**) Control (**B**) OTX2^CRISPR^. White arrows denote electroporated OTX2 positive cells. **C-H**. Representative dot plots showing the overlap between OTX2 (**C**), PAX6 (**E**), and THRBCRM2::GFP (**G**) with CAG::GFP or CAG::tdT in WT and OTX2^CRISPR^ dissociated retinas. Percentage of cells positive for each marker were normalized to the total number of electroporated cells detected with GFP or tdT. For OTX2 and PAX6 quantifications (**D, F**), p<0.0001. For THRBCRM2 comparison =0.0013. Error bars represent 95% confidence intervals, n=4 each WT and OTX2^CRISPR^. OR, outer retina; IR, inner retina.

The absence of OTX2-positive cells is more prominently observed at E10, when the WT retina is layered and the OTX2-positive PRs are concentrated in the ONL and BCs are located in the outer part on the INL (Suppl. Fig. 2C-E). Columns of GFP positive cells mutant for the OTX2 gene form gaps within the OTX2-positive populations in the ONL and INL (Suppl. Fig. 2D,E). These experiments suggest that Cas9 is able to disrupt OTX2 expression and leads to underrepresentation of PR and bipolar fates, in agreement with a previous study in mice (Nishida et al., 2003; Koike et al., 2007). To quantitate these effects and investigate the fate of OTX2 mutant cells, we turned to an *ex vivo* preparation at a later timepoint.

### Quantification shows a severe reduction in OTX2 along with an increase in PAX6 positive cells

To more precisely examine the role of OTX2 during PR genesis, we introduced the OTX2 CRISPR g2 into embryonic day 5 (E5) chicken retinas, a developmental time at which OTX2-positive RPCs are present and generating PRs and HCs. Retinas were electroporated *ex vivo* and cultured for two days. Immunofluorescence examination of retinas revealed a qualitative decrease in OTX2-positive cells in the electroporated population (arrows in Fig. 2A,B).

To quantify the effects presented in Fig. 2A,B, similar experiments were performed, followed by dissociation of the entire retina and quantification of the OTX2-positive cells using flow cytometry. In the OTX2^CRISPR^ mutants, an approximately 50% reduction in the percentage of OTX2-positive cells within the electroporated population was detected using two different OTX2 antibodies (Fig. 2C,D, Suppl. Table 1, Supp. Fig. 2F,G).

**Table. 1.** Antibodies used in the present study

| Antibodies | Source | Identifier |
| --- | --- | --- |
| Goat anti-OTX2 (1/500 from stock) | R&D Systems | Cat#AF1979; RRID: AB_1617988 |
| Mouse anti-PAX6 ( 1.125 ug/ml) | DSHB | RRID: AB_528427 |
| Mouse anti-TFAP2A (0.155 ug/ml) | DSHB | 3B5, RRID: AB_2313948 |
| Mouse anti-BRN3A (1/400 from stock) | Millipore | Mab1585, AB_94166 |
| Mouse anti-LIM1 (3.82 ug/ml) | DSHB | 4F2-C, RRID: AB_531784 |
| Mouse anti-ISL1 (0.205 ug/ml) | DSHB | 39.3F7, RRID: AB_1157901 |
| Rabbit anti-PH3 (0.5 ug/ml) | Millipore | Cat#06-570, RRID: AB_10582726 |
| Sheep anti-VSX2 (0.312 ug/ml) | Ex Alpha | Cat# x1180p, RRID: AB_2314191 |
| Chick anti-B-GAL (1/1000 from stock) | Abcam | Cat# ab37382, RRID: AB_307210 |
| Rabbit anti-GFP (1/500 from stock) | Invitrogen | Cat# A6455, RRID: AB_221570 |

In parallel, the percentage of PAX6-positive cells in the OTX2^CRISPR^ mutants was investigated, since previous experiments that used a mouse model of OTX2 mutation have shown an upregulation of this gene (Nishida et al., 2003). In OTX2^CRISPR^ mutants, an almost two-fold increase in the percentage of PAX6-positive cells was detected (Fig. 2E,F).

To test if PRs were also lost upon OTX2 mutagenesis, the number of cells that activate the THRBCRM2 cis-regulatory element was quantified. This element is associated with the THRB gene and is active in a subset of cone PRs in the chick retina (Emerson el al., 2013). OTX2^CRISPR^ mutant retinas had a significant reduction in PRs with THRBCRM2 activity (Fig. 2G,H). Thus, this assessment suggests that the OTX2^CRISPR^ mutation produced here in the chick leads to a robust downregulation of OTX2 protein and displays phenotypes similar to that previously reported in the mouse retina.

Quantification of all parameters reported above are detailed in Suppl. Table 1.

### The OTX2ECR2 reporter shows a change in cell fate in the OTX2^CRISPR^ cells

As OTX2 expression has been reported to be initiated in RPCs that predominantly generate cones and HCs, we examined these progenitors’ daughter cells to determine their fate upon OTX2 mutation. To do this, we used the previously reported OTX2ECR2 element (Suppl. Fig. 3A), which is active in OTX2 RPCs in the chicken retina (Emerson and Cepko, 2011). In WT retinas, GFP expression driven by the enhancer’s activity is observed predominately in the outer retina/PR layer, while fewer cells, showing less intense GFP expression, in the inner retina (Suppl Fig. 3B-C). The enhancer’s activity co-localizes primarily with OTX2-positive retinal cells electroporated at E5 and cultured for 2 days consistent with previous observations (Emerson and Cepko, 2011). *In vivo* experiments have demonstrated that OTX2ECR2 drove reporter activity in both PRs and LHX1-positive HCs and very rarely targeted BRN3A-positive RGCs (Emerson and Cepko, 2011).

**Figure 3.**
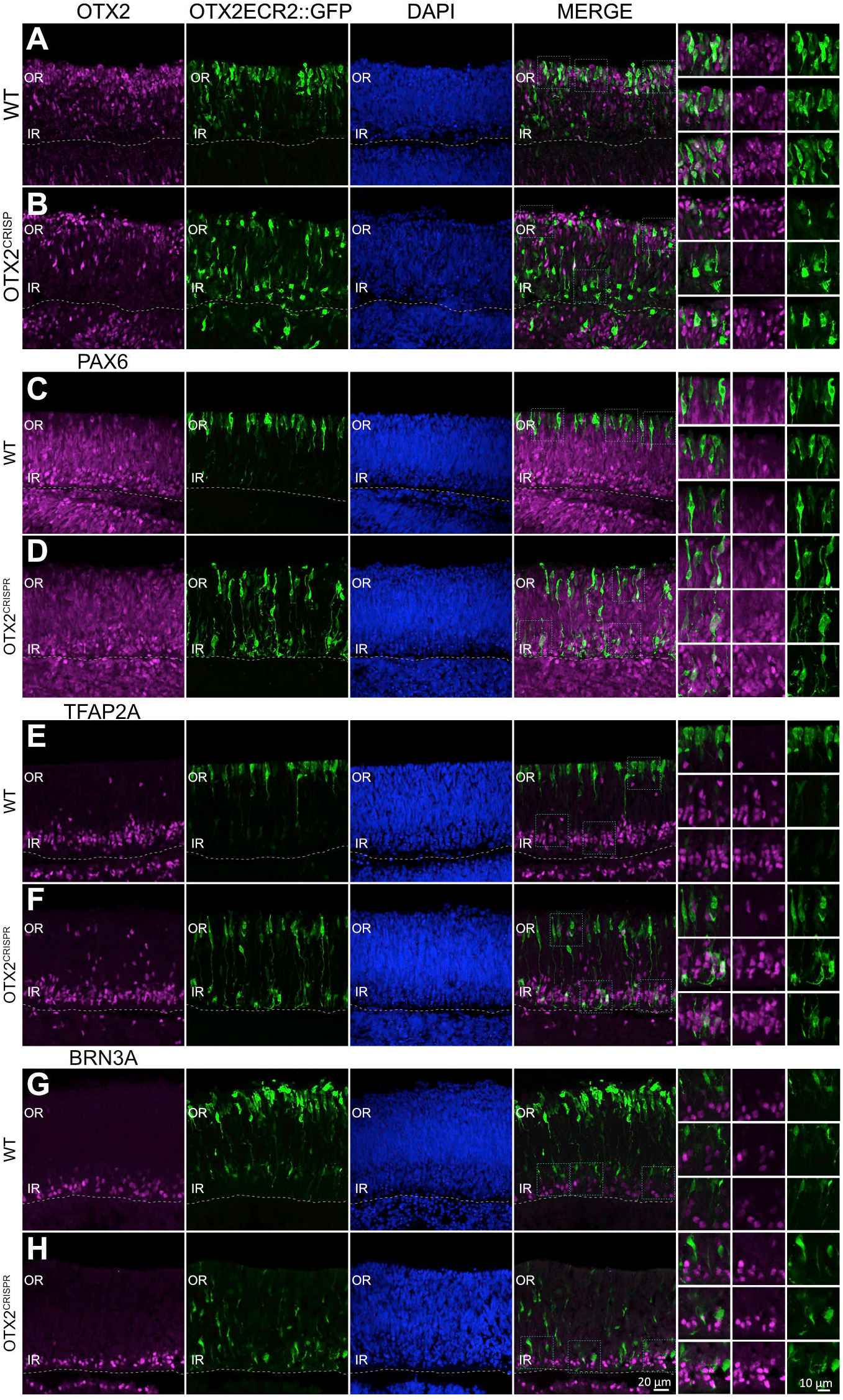
Immunofluorescence characterization of OTX2ECR2 reporter in WT and OTX2^CRISPR^ retinas. Retinas were co-electroporated at E5 with OTX2ECR2::GFP and Control or OTX2 CRISPR plasmids and analyzed after 48 hours. Boxed regions in the Merge column are shown at higher magnification in the last 3 columns. **A-H**. OTX2, PAX6, TFAP2A and BRN3A expression in WT (**A, C, E, G**) and OTX2^CRISPR^ (**B, D, E, H**) retinas.

To confirm previous estimates of the activity of the OTX2ECR2 element in OTX2-positive cells, a quantitative flow cytometry analysis was performed. 73% (c.i. ± 2.4%) of GFP-positive cells were also OTX2-positive. In contrast, in the OTX2^CRISPR^ mutants the activity of the enhancer overlaps with the protein in only 22% (c.i. ±5.9%) of the cells (Suppl. Fig. 3D,E, Suppl. Table 1). In addition, there is a robust increase in both the number and the intensity of the signal in the GFP-only population, suggesting that these are the newly generated, mutant cells (Suppl. Fig. 3C, D).

In conclusion, characterization of the OTX2^CRISPR^ retinas shows a qualitative shift in the cell populations labeled by the OTX2ECR2 reporter – a dramatic reduction in the PR population as well as the formation of a new population in the inner retina, with cells having strong GFP expression driven by OTX2ECR2 (Suppl. Fig. 3).

### Characterization of fate changes in the OTX2^CRISPR^ mutants

In the OTX2^CRISPR^ mutants, the few remaining OTX2ECR2::GFP-positive PRs are OTX2-positive, suggesting that these cells likely still retained at least one functional OTX2 allele, that they were not efficiently triggered for Cas9 cleavage, or that the induced mutation was hypomorphic. (Fig. 3B,D,F). Since the newly formed OTX2ECR2 population is located in the inner retina, markers for RGCs, ACs or HCs were used to initially characterize these cells.

Based on the increase in the number of PAX6-positive cells in the OTX2^CRISPR^ mutants, the overlap of the newly formed inner retina population with PAX6 was analyzed both qualitatively and quantitatively. Indeed, OTX2ECR2::GFP-positive cells in the wildtype are Pax6-negative while those in the inner retina of the OTX2^CRISPR^ mutants are PAX6-positive (Fig. 3C,D). In addition, an increase in the percentage of PAX6-positive cells in the OTX2ECR2 population from 0.3% (c.i. ±0.9%) in the WT retinas to 8.4% (c.i. ±4.3%) in the OTX2 mutants was observed using flow cytometry, with a p=0.042, n=4, when a two tailed student’s ttest was performed.

Similar analysis was performed using TFAP2A as marker for ACs and HCs (Fig. 3E,F). Quantitative analysis yielded a mean of 4.4% (c.i. ±1.9%) overlap between the marker and OTX2ECR2 cells in the WT retinas and 8.3% (c.i. ±2.8%) in the OTX2^CRISPR^ mutants, with a p=0.069, n=4, when a two tailed student’s ttest was applied. Very few OTX2ECR2 cells appear to be positive for the RGC marker POU4F1 (BRN3A) in the OTX2^CRISPR^ mutants when confocal micrographs were assessed qualitatively (Fig. 3G,H).

### Droplet based single cell RNA sequencing analysis of the OTX2ECR2-positive cells

To investigate the fate of OTX2ECR2 population in the OTX2^CRISPR^ retinas at a higher resolution, we collected OTX2ECR2::GFP-positive cells from both mutant and WT retinas based on their GFP florescence (Suppl. Fig. 3F) after *ex vivo* electroporation at E5 and two days in culture. Cell suspensions were then processed for single cell library preparation using a 10X Genomics Chromium Single Cell 3’ platform (Zheng et al., 2017) and sequenced on an Illumina HiSeq 4000 sequencer. Prior to retinal dissociation, the embryos were genotyped for sex determination (Methods) and cells from four retinas of embryos PCR-identified as female were pooled per sample. For each of the mutant retinas included in the sample, the contralateral eye was used as WT control to limit the variability across individual animals.

In total, 5,122 cells were recovered from the WT sample and sequencing yielded an average of 65,555 reads per cell, with a total number of 17,485 genes detected and a median of 1,293 genes per cell. From the OTX2^CRISPR^ sample, 4,940 cells were recovered, sequencing yielded an average of 66,303 reads per cells with a total of 17,351 genes detected and a median of 1,254 genes per cell.

To classify the cells in the OTX2ECR2 lineage in the WT scenario, filtered cells were run through a linear dimensional reduction algorithm using highly variable genes in Seurat (Methods), computing 30 principal components (PCs). Mitochondrial, ribosomal and cell cycle genes were regressed from the analysis as discussed in the methods section (Suppl. Table 2). All computed PCs were analyzed with the JackStraw procedure (Macosko et al., 2015) and 22 were processed further for cluster analysis. The FindClusters function was used in Seurat to compute the clusters, which were then visualized with the t-distributed stochastic neighbor embedding (tSNE) algorithm.

Ten different clusters were identified in the WT OTX2ECR2 lineage based on previously characterized markers (Fig. 4A). As expected, not all clusters presented high levels of OTX2, supporting the previous data on the characterization of the OTX2ECR2 enhancer (Emerson and Cepko, 2011). Namely, clusters 1, 2, 3, 4 and 5 show high levels of OTX2 in the majority of cells, whereas clusters 0, 6, 7, 8 and 9 show only sparse cells positive for the marker, with minimal expression levels (Fig. 4C,D). These could represent cells that either have a previous history of OTX2 expression or an inappropriate activation of the OTX2ECR2 element.

**Figure 4.**
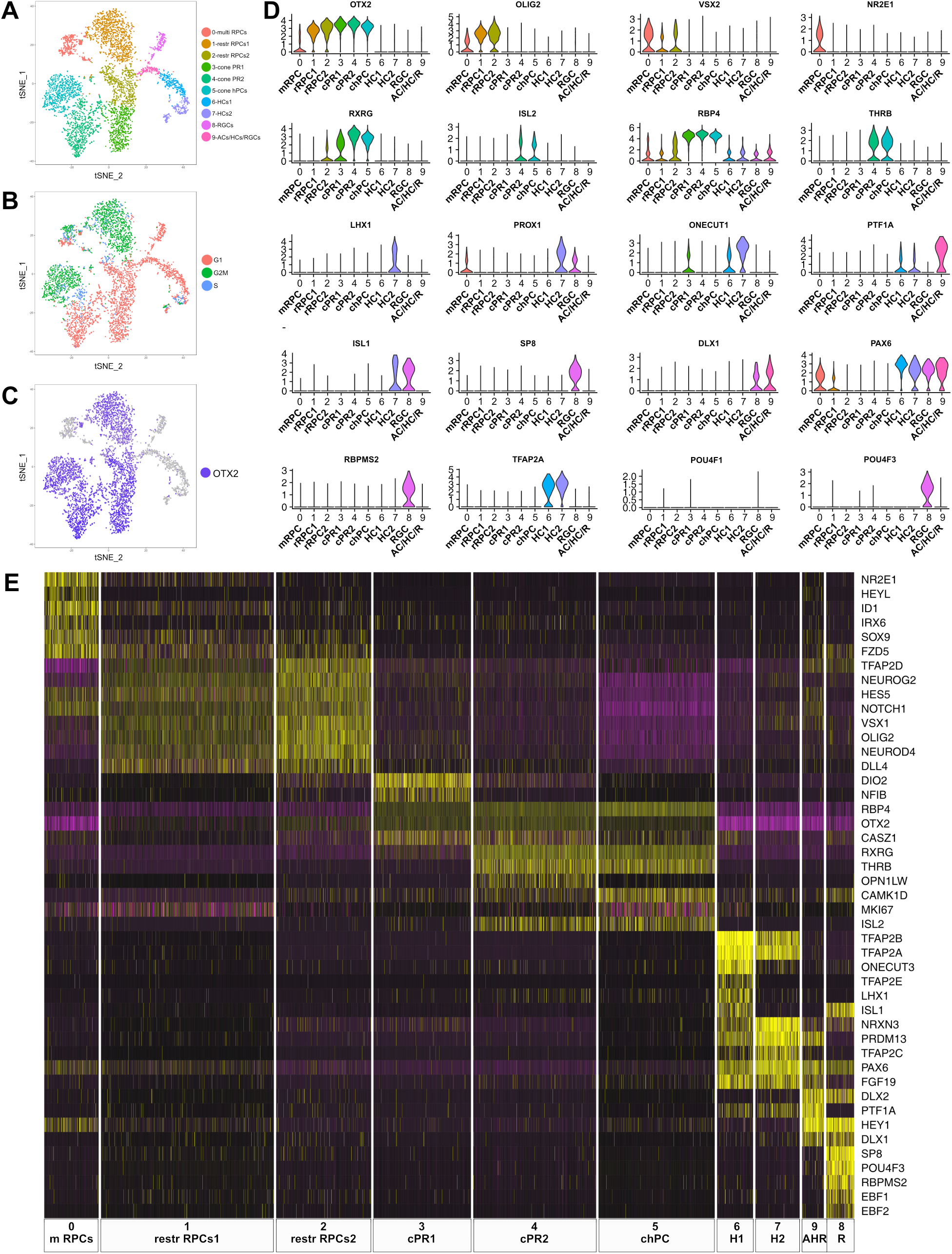
Single cell analysis of WT OTX2ECR2::GFP cells. Retinas were electroporated at E5 and dissociated after 48hrs. **A**. TSNE plots of the 10 clusters generated by the unsupervised algorithm Seurat, based on the gene expression of each cell analyzed. **B**. TSNE plots of the clusters showing their cell cycle state, G1, G2M or S-phase. **C**. OTX2 expression in the cells analyzed (purple). **D**. Violin plots showing the distribution of different markers in the 10 clusters. **E**. Heatmap of differentially expressed genes across the 10 clusters, where purple represents low gene expression and yellow represents high expression.

Two OTX2 and OLIG2 positive clusters were identified as containing restricted RPCs, cluster 1-restr RPCs1 with cells undergoing G2M or S-phase, (1121 cells), and cluster 2-rest RPCs2 with cells in G1 phase (616 cells) (Fig. 4A, B, D). Based on gene expression profiles, these are likely to represent the same cell state in different stages of the cell cycle. 353 cells were assigned to be multipotent retinal progenitor cells (cluster 0-multi RPCs) based on their high expression of VSX2, NR2E1, ID1 and SOX9.

Two different clusters – 3-cone PRs1 (632 cells) and 4-cone PRs2 (797 cells) are assigned to be PRs, which are presumably cone PRs based on the early time point of examination as well as the markers expressed. We have categorized cluster 5 as cone homotypical progenitor cells (chPCs) (754 cells) based on their very similar gene expression profiles to cluster 4, with the addition of cell cycle markers. All three clusters present high levels of RBP4, with expression of THRB, RXRG, and ISL2 in clusters 4-cone PRs2 and 5-chPCs and lower levels of RXRG in 3-cone PRs1.

Cluster 8-RGCs was identified as containing RGCs (181 cells), based on its high expression of POU4F2 and POU4F3, RBPMS2, as well as members of the EBF family. Surprisingly, POU4F1, which is present in a large fraction of RGCs was only found in a minority of the cells in this cluster (Liu et al., 2000). At least two clusters were identified to be horizontal cells (HCs), cluster 6 and 7 (278 and 233 cells, respectively), both containing cells that express PROX1, ONECUT1, TFAP2A and TFAP2B. The 141 cells in cluster 9-ACs/HCs/RGCs express the horizontal/amacrine cell (AC) marker PTF1A almost exclusively, but also members of the DLX family of transcription factors, known to be characteristic for murine ACs and RGCs (Melo et al., 2003). However, due to the similarity in their transcription programs and to the lack of specific markers for ACs at this developmental time point, the discrimination between the three classes is difficult to make (Fig. 4E, Suppl. Fig. 7, Suppl. Tabel 3), (Clark et al., 2018 BioRxiv).

### OTX2^CRISPR^ mutants lack all PRs clusters and show increase numbers of RGCs and HCs

To compare the single cell transcriptomes from both WT and OTX2^CRISPR^ samples, a similar cluster analysis based on the principal component was run with the two datasets combined (Fig. 5A, B). From the 40 total PCs computed, 33 were processed further. All 10 clusters assigned when WT cells were analyzed individually were present after this analysis, with minimal differences in the number of cells per cluster

**Figure 5.**
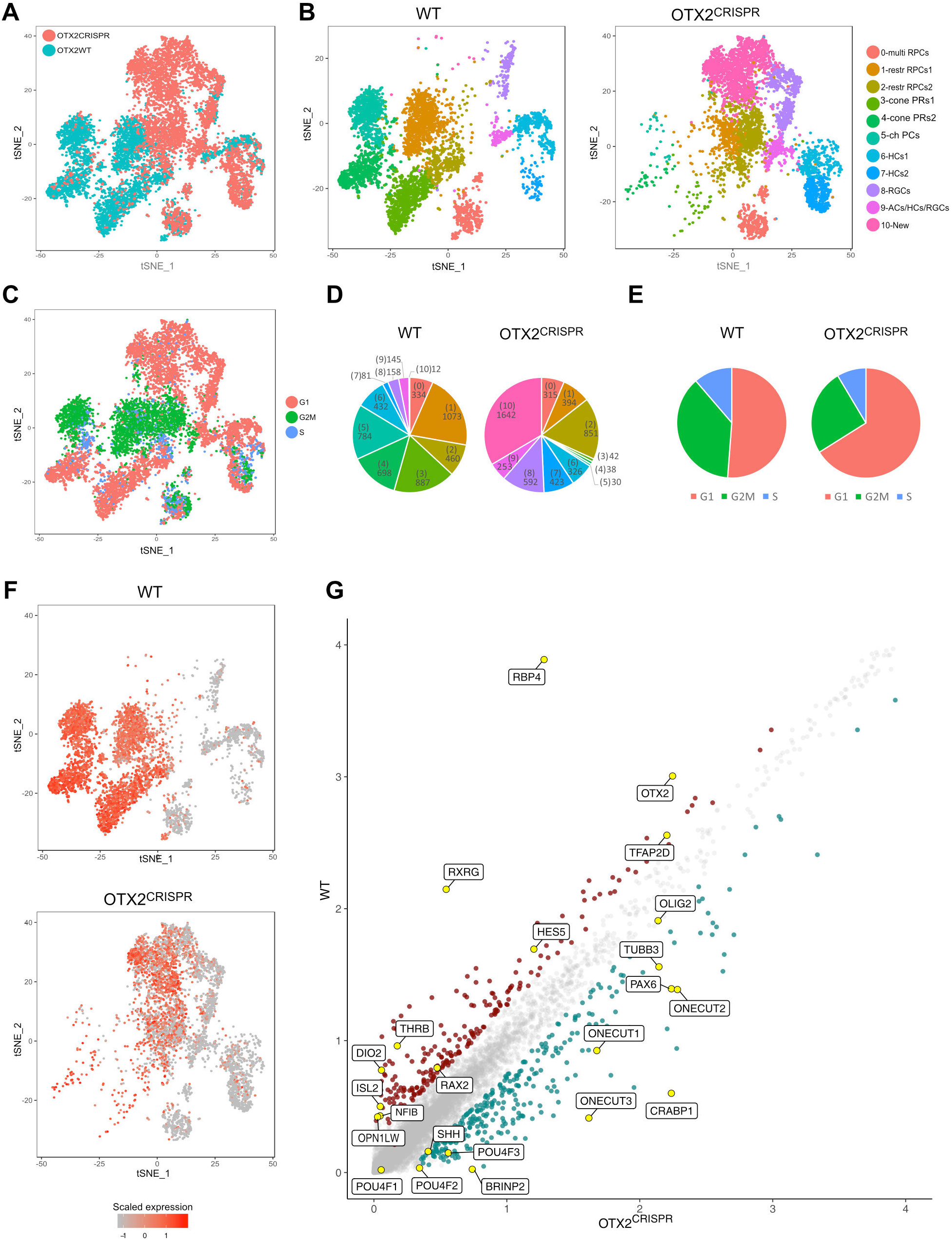
Combined single cell analysis from WT and OTX2^CRISPR^ retinas labeled by OTX2ECR2::GFP reporter. TSNE plots of the two datasets analyzed simultaneously (**A**) and labeled for their cell cycle signature (**C**). **B**. TSNE plots of the 11 clusters determined by Seurat – 0-multipotent RPCs, 1 and 2 restricted RPCs, 3-5 PRs, 6 and 7 two HCs clusters, 8-RGCs, 9-ACs/HCs/RGCs and 10-New in the WT (left) and mutant (right) samples. **D**. Pie charts showing the number of cells found in each cluster of the WT and mutant samples, cluster numbers are in parentheses, **E**. number of cells found in different cell cycle phases. **F**. Heatmap of OTX2 expression across the two datasets. **G**. Differentially expressed genes in the WT and OTX2^CRISPR^ cells showing average reads per cell.

To estimate the extent of OTX2 mutagenesis, the sequencing reads around the Cas9 target site were analyzed in both the mutant and WT samples (Suppl. Fig. 4). Examination of the sequencing reads using the Integrative Genomics Viewer – IGV (Robinson et al., 2011) shows a wide degree of mutation present at that location (Suppl. Fig. 4B). While the majority of the reads are located at the 3’ end of the gene (Suppl. Fig. 4A), the targeted second coding exon in the OTX2^CRISPR^ sample has a diverse set of mutations, presumably introduced through the NHEJ repair pathway.

Widespread changes in cell cycle states and the distribution of cells of the OTX2^CRISPR^ sample relative to the WT clusters were observed (Fig. 5C, E and Suppl. Fig. 4C-D). All three PR clusters were massively reduced in the OTX2^CRISPR^ sample to approximately 5% of their levels found in WT (Fig. 5B, D). OTX2 mRNA expression was dramatically reduced in the OTX2^CRISPR^ sample but not completely lost in the mutant cells, whereas the majority of the PR markers were severely decreased – THRB, RXRG, RBP4, ISL2, OTX5 (Suppl. Fig. 5D-G, Suppl. Fig. 7). The cells that continue to express OTX2 mRNA likely have mutations in the OTX2 gene that lead to the loss of functional OTX2 protein. The loss of PR markers is also observed when data is analyzed in a “bulk-RNAseq”-like fashion, with all transcripts analyzed concomitantly (Fig. 5G, Suppl. Fig. 4).

The restricted RPC populations together were reduced approximately 19% in the OTX2^CRISPR^ condition, most having G2/M states. (Fig. 5E, Suppl. Fig. 4D).

In the cell clusters that were not actively expressing OTX2, specific cell populations were increased while others were largely unaltered. The multipotent RPCs found in cluster 0 were unaffected, as expected, given the lack of OTX2 expression in these cells – 334 cells in WT, 315 in the OTX2^CRISPR^ (Fig. 5D-E, Suppl. Fig. 7). However, a moderate increase was noticed in cluster 9 – ACs/HCs/RGCs in the OTX2 mutants, from 145 cells in WT to 253 mutant cells (Fig. 5D). Examination of clusters with more terminal fates provided more insight into the consequences of OTX2 mutation. The number of RGCs in the OTX2^CRISPR^ sample increased by approximately 4-fold, from 158 cells in the WT retinas to 592 in the mutants (Fig. 5D, Suppl. Fig. 7). The number of BRN3A (POU4F1) positive cells was not increased in the OTX2^CRISPR^ cells, confirming the characterization of the mutant population using immunohistochemistry (Fig. 3G-H). However, the other two members of the POU4F family, POU4F2 and POU4F3, were dramatically increased in the OTX2^CRISPR^ cells, suggesting that only a certain subpopulation of RGCs was formed upon OTX2 mutation.

The combined analysis revealed that the two HC clusters also changed in specific ways. Strikingly, the LHX1-positive HCs (cluster 7 – HCs2) increased by 5-fold in the OTX2^CRISPR^ sample. For cluster 6 HCs1, a further analysis of both datasets combined revealed that some cells are also LHX1-positive. Subclustering of cluster 6 revealed that as expected (Fischer et al., 2007), there is a population of cells expressing mostly ISL1, subcluster 6-1 (Suppl. Fig. 6), while a different subcluster consists of the LHX1-positive cells, cluster 6-2, and a third contains cells negative for both markers, 6-0. The overall number of cells in cluster 6 – HCs1 decreased in size by approximately 25%, from 432 cells in the WT to 326 cells in the mutants (Fig. 5D, Suppl. Fig. 6, Suppl. Fig. 7). However, the only subcluster that showed an increase in cell number in the OTX2^CRISPR^ cells, from 12 in the WT to 145, was the LHX1 positive subcluster 6-2, following the same trend as cluster 7 (Suppl. Fig. 6).

**Figure 6.**
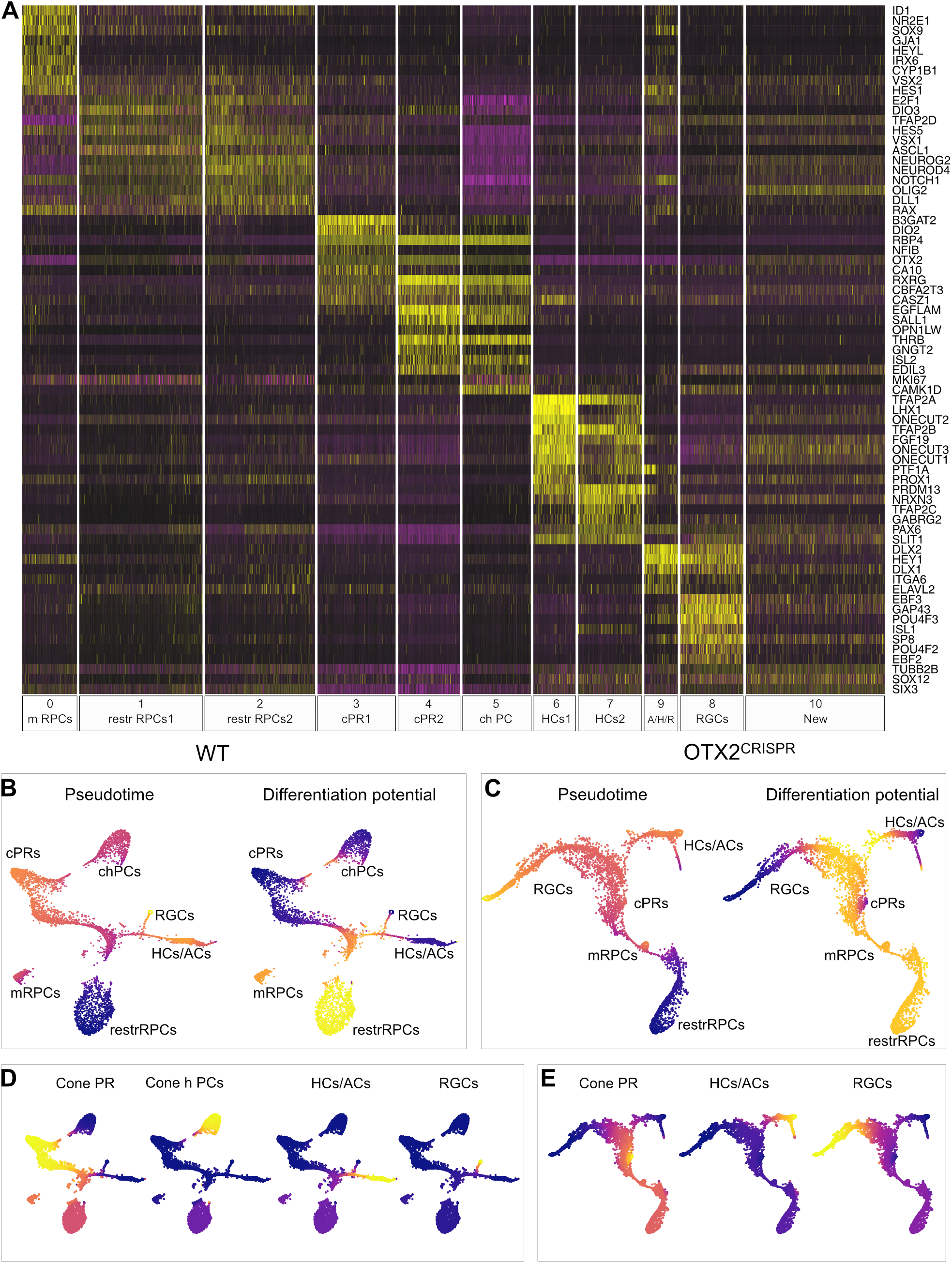
Differential expression of marker genes across clusters. **A**. Heatmap of the differentially expressed genes across the 11 clusters of both WT and mutant datasets. Purple represents low gene expression and yellow represents high expression. **B-E**. Pseudotime and differentiation potential model for WT (**B, D**) and OTX2^CRISPR^ (**C, E**). Pseudotime represents a model of cell ordering across all lineages relative to the start cell, input as the restRPC population. In the differentiation potential view, the cold, dark blue shades represent final states, while warmer colors show higher differentiation potential, chPCs, cone homotypic progenitor cells.

**Figure 7.**
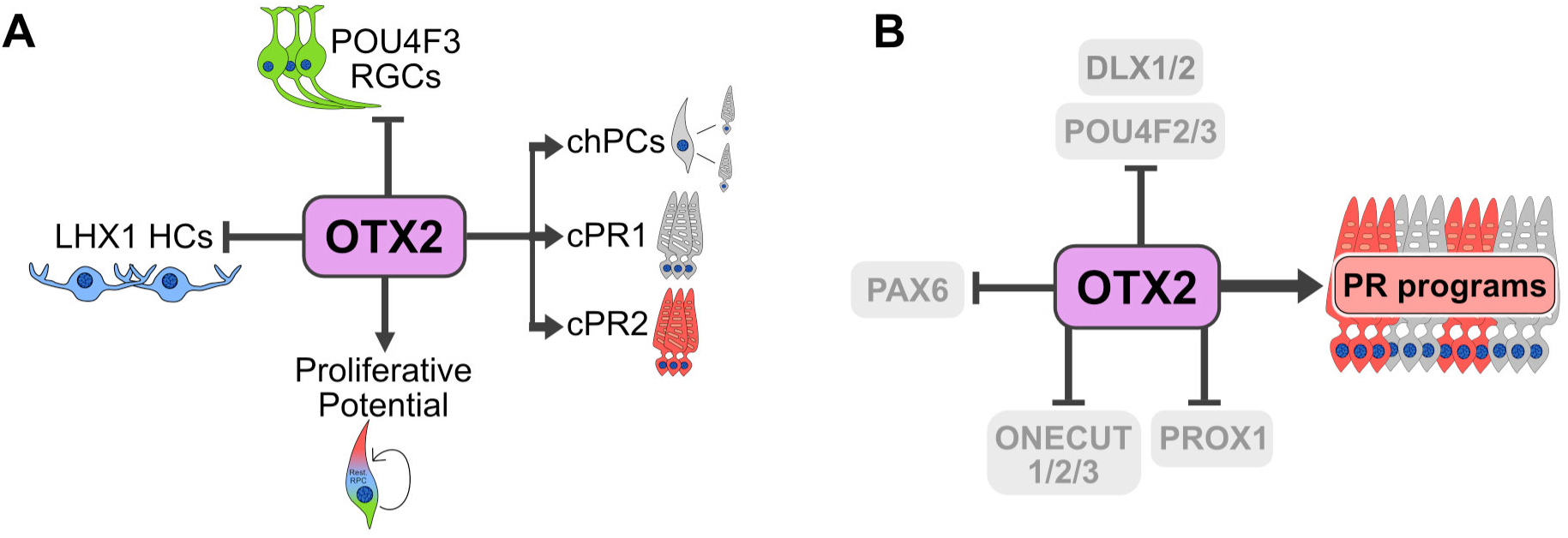
Contribution of OTX2 during the cell fate specification in the developing chick retina. **A**. OTX2 is necessary for the generation of two PRs types, as well as one type of cone homotypic progenitor cell (chPCs) and ensures the division potential of restricted progenitor cells, while repressing the generation of subtypes of RGCs and HCs. **B**. OTX2 activates genes required for formation of PRs, inhibiting regulators of RGC (POU4F2/3, DLX1/2), HC (ONECUT, PROX1) and PAX6 expression.

Taken together, there appears to be an increase in a number of specific cell fates, including LHX1-positive HCs and POU4F2/POU4F3 RGCs, while other populations such as multipotent RPCs, ISL1-positive HCs, and POU4F1 RGCs are relatively unchanged. The increase in HC and RGC fates is consistent with the increased INL populations observed in the retina (Suppl. Fig 3C).

In agreement with the immunohistochemical characterization, PAX6 was expressed in a large number of cells in the mutants compared to the WT. Moreover, a concomitant double analysis of OTX2 and PAX6 transcripts at the single cell level shows virtually no overlap between the two markers in WT and minimal overlap (green dots in Suppl. Fig. 5A) in the OTX2^CRISPR^ retinas, suggesting a negative correlation between these two factors. A derepression of PAX6 was identified even within the restricted RPC populations, suggesting that it was a primary transcriptional response to OTX2 loss, and not just a secondary effect of cell fate change. The cells expressing both OTX2 and PAX6 in the mutant retinas could potentially represent cells that carry mutant OTX2 transcripts. In addition, all three members of the ONECUT family, ONECUT1-3, had increased expression after OTX2 mutation, suggesting a potential inhibition of these factors by OTX2 in WT conditions (Suppl. Fig. 5B).

### One third of OTX2 mutant cells form a unique cell class

As mentioned previously, the combined analysis of the WT and mutant cells revealed a new cluster, cluster “New”. Initial analysis of the markers suggested no connection to a known retinal cell class at any time point, and certainly at the time point of this analysis. These cells are in G1/G0, according to the analysis of the cell cycle genes (Suppl. Fig. 4D) and are completely lacking the cell proliferation marker MKI67. The cycling-dependent kinase inhibitor – CDKN1A, a gene associated with cell cycle exit (Bunz et al., 1998), is highly expressed in cluster “New” and is minimally expressed in the WT sample. In addition, amongst the highly expressed genes in the “New” cluster is BRINP2 (BMP/retinoic acid inducible neural-specific 2), which has also previously been shown to promote cell cycle exit (Kawano et al., 2004), and CRABP1 (cellular retinoic acid binding protein 1), suggesting a potential aberrant retinoic acid signaling in these cells (Suppl. Fig. 7).

A subsequent analysis of cluster “New” revealed that these cells can be sub-classified into three clusters (10-0 to 10-3) (Suppl. Fig. 5E). Subcluster 10-1 represents 559 cells in the mutant data set compared to 9 WT cells, which express high levels of RAX, NOTCH1, NEUROG1, HES5, NEUROD4 and HES4, all markers that suggest a potential progenitor signature (Suppl. Fig. 5H). Subcluster 10-0 is formed by 554 OTX2^CRISPR^ cells and 4 OTX2 WT cells that express markers that implicate a potential RGC signature, such as EBF3, GAP43, POU4F2 or PCHD9, in addition to a few markers previously associated with HCs and ACs, like MEGF10 and MEGF11 (Suppl. Fig. 5G). Lastly, subcluster 10-2 comprises 539 mutant cells and 9 WT cells. These cells have the highest expression of OTX2 amongst the three subclusters in cluster “New”, and express additional PR markers – RBP4, RXRG, PDE6G, PRDM1, or markers that have recently been associated with a PR signature – CBFA2T3, CAMK1D and CLSTN1 (Suppl. Fig. 5F) (Buenaventura et al., 2018).

To further investigate the composition of cluster “New”, we followed a different bioinformatic approach to look at potential single cell trajectories ordered by pseudotime with the purpose of modelling the temporal profile of this cluster “New” in the cell differentiation process. The analysis of the WT-only dataset using the Palantir program (Setty et al., 2018 Biorxiv) confirmed the assigned terminal cell fates previously obtained using Seurat – PRs, ACs/HCs and RGCs (Fig. 6B, D). Similar analysis looked at cells collected from the mutant retinas and identified only two main terminal fates branching from the restricted RPCs – ACs/HCs and RGCs, with a minimal representation of cone PRs (Fig. 6C, E). Distribution of markers characteristic for cluster 10 placed it in between the restricted progenitors and the final fate branches, further suggesting the transitory state of these cells (Suppl. Fig. 6D). Moreover, most of the cells in cluster 10 were emerging towards the RGC branch.

## Discussion

The utility of the CRISPR/Cas9 system in the induction of targeted mutagenesis has been firmly established across multiple paradigms, not only facilitating the study of somatic mutations at any given time during development, but also allowing these studies in less genetically-tractable model organisms, like the chicken (Gandhi et al., 2017). Furthermore, the ability to introduce mutations in specific tissues at particular timepoints has proven particularly useful for developmental studies, allowing for a more precise evaluation of genes that have a role in multiple developmental time windows. In addition, the recent technological advances for the generation of single cell gene expression profiles allows for analysis of rapidly dividing and differentiating cells at a single cell resolution (Carter et al., 2018; Zeisel et al., 2018). The current study illustrates the power of using these methods in combination to evaluate gene function in the context of a developing tissue. This has enabled an extensive characterization of OTX2 mutant cells in the developing retina, supporting the active role of OTX2 homeodomain TF in the commitment of RPCs to a PR fate as well as its role in repressing the generation of specific subtypes of RGCs and HCs.

### Timing and CRISPR effectiveness of the OTX2 ablation

Introduction of OTX2^CRISPR^ at multiple timepoints allowed for identification of previously described phenotypes for OTX2, including effects on eye morphology, RPE pigmentation/gene expression, PR and BC formation, and repression of PAX6 expression. Moreover, this current mutation model resembles the human mutations of OTX2 that lead to different degrees of microphthalmia and anophthalmia more than any other reported model of OTX2 ablation. These malformations were reported to occur as an effect of *de novo* nonsense mutations that lead to early termination of OTX2 transcription and either lead to degradation of the transcript by nonsense-mediated RNA decay or to dysfunctional OTX2 protein due to defects in the transactivation domain (Boyl et al., 2001; Gat-Yablonski 2011). The efficacy and versatility of this CRISPR electroporation approach is in line with previous studies of the chick neural tube and the mouse retina and cortex (Timmer et al., 2001). The quantitative analysis at E5 shows that OTX2 protein-positive cells were significantly reduced by OTX2^CRISPR^, and the analysis of the OTX2 mRNA reads in the single cell fashion shows that mutation induction was highly efficient. The biological effects gauged using PR reporters and cell type-specific markers were confirmed by the single cell analysis.

One important consideration of the CRISPR technique is the possibility of off-target effect of Cas9 activity, which can be a potential source of multiple-gene mutants. In the present study, the most expected off-targets of OTX2 g2 are OTX1 and OTX5 (CRX), both members of the same family. CRX has been shown to be downstream of OTX2, and their DNA sequences are very distinct, limiting the possibility of representing a target for OTX2 guide 2. OTX1 is associated with optic stalk development and is expressed at insignificant levels in the developing neuronal retina at E5, the timepoint where the single cell profiling was carried out. Investigation of the OTX1 transcripts in single cell transcriptomes shows that this gene is expressed more in the OTX2^CRISPR^ cells compared to WT, and, therefore suggests a potential compensation loop after OTX2 ablation. In addition, despite its low sensitivity, an enzyme mismatch cleavage assay confirmed that OTX1 is not a target of Cas9 OTX2 guide2 (data not shown).

### Cell type-specific effects in response to loss of OTX2

Previous analysis of OTX2 function in the formation of the mouse retina has concluded that conditional loss of OTX2 in postmitotic PRs or throughout the retina using an early neural retina driver leads to loss of PRs, HCs, and BCs and to a concomitant increase in ACs. While the differences observed in this study could be due to a mouse/chick distinction, we favor the explanation that three experimental aspects in this study allowed for novel findings: 1) short-term assessment after OTX2 mutation, 2) a targeted cell population that includes OTX2-positive RPCs and 3) single cell analysis to sensitively identify cell populations. The three previous developmental studies examined the effects of OTX2 mutation beginning either in post-mitotic PRs, or in the early retina prior to postmitotic cell formation and possibly at a timepoint when OTX2 is still critical for eye specification (Nishida et al., 2003; Sato et al., 2007; Omori et al., 2011). In addition, in two of the previous cases, the evaluation of cell fate largely occurred in the adult, possibly precluding the detection of the cell cycle and cell fate changes observed in this study. However, several of the gene regulatory changes identified here associated with HC and RGC formation were identified in the previous study that examined the OTX2 CRX conditional knockout at postnatal day 1 (Omori et al., 2011). These include upregulation of PROX1, associated with HCs, BPs, and aII ACs, and of POU4F2 (BRN3B), expressed in RGCs. In addition, DLX1 and 2 were strongly upregulated. These gene expression changes are consistent with the ones we observed in the chick retina and may indicate a similar increase in the mouse gene regulatory networks associated with HCs and RGCs.

### Photoreceptors

As previously observed in the mouse retina, PRs were the cell type most obviously lost in all OTX2 loss-of-function conditions. In the wildtype cells labeled by OTX2ECR2, there are two distinct postmitotic cone clusters present in the retina and both appear to be missing as a consequence of OTX2 mutation. All genes associated with PRs were drastically downregulated and their absence in any of the cell clusters in the OTX2 mutants suggests these genes are directly or indirectly downstream of OTX2. This is congruent with the proposed role for OTX2 as a pivotal upstream component of the PR gene regulatory network.

### Horizontal Cells

The mouse retina has been characterized as only possessing one type of HC, the Type 1 distinguished by LHX1 (LIM1) expression, and it was reported to be missing in the OTX2 mutant (Sato et al., 2007). In this study, we find that Type 1 HCs are dramatically increased suggesting that OTX2 is not required for this cell type in the chick retina, at least during this temporal window. In both species, at least some HCs and cone PRs are born from the same RPCs and in the chicken these have been defined by the activity of the THRBCRM1 element or OLIG2 (Hafler et al., 2012; Emerson et al., 2013). These cells were likely targeted at E5 to generate some of the cone and HCs profiled. Interestingly, lineage tracing experiments of the THRBCRM1 population *in vivo* has determined that these cells preferentially generate the Type 1 HCs over the Types 2, 3, and 4 defined by ISL1 expression (Schick et al. 2019, BioRxiv). This suggests that one of the functions of OTX2 in the THRBCRM1 population or other OTX2 RPCs is to specifically repress the Type 1 HC fate in PR progeny. It will be interesting to investigate at what node in the HC gene regulatory network OTX2 acts to repress this fate. One possible set of targets are the three members of the ONECUT family of genes. Previous analysis has determined that these genes are necessary for the formation of LHX1-positive HCs in the mouse, that misexpression of ONECUT1 alone is sufficient to induce these cells, and that chicken LHX1-positive cells express high levels of ONECUT1 (Wu et al., 2013; Sapkota et al., 2014; Emerson et al., 2013; Buenaventura et al., 2018). The large increase in ONECUT1/2/3-positive cells we observed in response to loss of OTX2 suggests that OTX2 normally represses these genes, perhaps keeping the LHX1 HC program off. Intriguingly, an LHX1 reporter in zebrafish drives reporter expression not only in HCs, but also in PRs (Boije et al., 2015). Though evidence for endogenous expression of LHX1 is lacking, including in the present single cell analysis, this could suggest that LHX1 expression is primed within these RPCs.

### Retinal Ganglion Cells

Retinal Ganglion Cells make up a minority of the cells found in the OTX2ECR2 population, but those present are primarily POU4F2 and POU4F3-positive cells and not POU4F1. As with HCs, the same timing and model differences could account for the discrepancy between these results and those of the previous mouse studies. In our recent analysis of a lineage trace analysis of THRBCRM1 cells, we observed that there is a very small number of RGCs formed from this population. Though the numbers are limited, it is intriguing that POU4F1-positive RGCs, like Type 2-4 HCs, are underrepresented in this population (Schick biorxiv). This may suggest that, like HCs, the specific induction of POU4F2 and POU4F3-positive cells is related to the limited cell fate choices of THRBRM1-positive cells. We observed that RGC genes were highly overrepresented in several clusters of the OTX2ECR2 population. These include the increased expression of the DLX1 and DLX2 genes that have been identified as primary regulators of the RGC fate (Zhang et al., 2017) in cell populations that normally represented RGC formation. In addition, the expression of the RGC-associated genes EBF3, GAP43, and RBPMS2 was consistently high along the course of cells from the focal point where HCs/ACs/RGCs form, suggesting that these cells are formed through some normal RGC genesis pathway. In the New cluster of OTX2 mutants, there are a number of cells that express these factors, though not as consistently as in WT clusters. We speculate that these cells have derepressed RGC transcriptional programs, but due to the timing of their OTX2 deletion, they either possess remnants of the OTX2 program that cause them to be clustered apart from the other developing RGCs, or they are led through an alternative pathway to the activation of RGC-associated gene regulatory networks. The earliest marker associated with RGC genesis, ATOH7, is also upregulated in the OTX2^CRISPR^ data set (data not shown). However, technical limitations do not permit the visualization of this marker at single cell level. One of the potential explanations for why the ATOH7 transcripts were not present in the single cell analysis might be that this marker aligned to more than one locus and the ATOH7 transcripts were filtered out as a result.

### Restricted RPC populations

Two clusters of restricted RPCs labeled by the OTX2ECR2 element were classified in the WT dataset. Based on their gene expression similarities, these two clusters seem likely to be the same cell type but are split between the cells in G1 and in G2-M. We are denoting these as restricted RPCs based on four criteria – 1) their clear segregation from multipotent RPCs, 2) their expression of cell cycle genes (at least for cluster 1), 3) their expression of genes noted in other restricted progenitors such as OLIG2 and ASCL1, and 4) a reduced expression of multipotent RPC genes such as PAX6, NR2E1, RAX and VSX2. These two populations appear to be distinct from the THRBCRM1 population, as neither of these populations express THRB and there is more VSX2 expression than would be expected in the THRBCRM1 population (Buenaventura et al., 2018). However, a flow cytometry experiment using co-electroporation of the THRBRM1 and OTX2ECR2 reporters confirmed that more than 50% or OTX2ECR2 positive cells were also positive for ThrbCRM1 at early timepoints in RPCs (data not shown). This suggests that many of the postmitotic cells in the single cell datasets are derived from ThrbCRM1 RPCs. The cell types that are formed from these other OTX2-positive RPC populations and whether these cells are restricted in their fate potential or not for this timepoint is unknown. In the OTX2^CRISPR^ condition, the relative proportion of the restricted RPC clusters change, with an increase in the G2 cluster, a decrease in the G1 cluster and a rightward shift in the cluster location, suggesting a shift in the gene expression profile of these cells.

### Homotypic RPC population

There are at least two other populations that, despite not being defined as RPCs, have robust expression of cell cycle genes and are clearly segregated from multipotent RPCs, which we categorize as homotypic RPCs. Homotypic PR, HC, and BC populations have been most readily identified in fish, with some evidence of their existence in chick (Rompani and Cepko 2007; Vitorino et al., 2009; Suzuki et al., 2013). One of these clusters, 5-chPCs expresses THRB and is extremely similar to cluster 4-cone PRs2, except that it is a cycling population and does not express OPN1LW. We hypothesize that the 5-chPC cells represent a homotypic RPC population that will generate long wave opsin positive cells found in cluster 4. In response to the OTX2^CRISPR^ introduction, this RPC population is almost entirely missing. In addition, we observe a subpopulation of cycling HCs– 7-HCs2, consistent with the previously identified population of horizontal RPCs (Godinho et al., 2007).

### New cluster in OTX2 Mutants

The new cluster 10 found in the OTX2 mutants is likely composed of cells that were destined to be PRs. The similar numbers of cells in the PR group and the New cluster, as well as the presence of some cells, especially in the top third of the cluster with a PR signature (Fig. 9), supports this idea. Interestingly, the vast majority of these cells are in G1, suggesting that the population we are denoting as cone homotypic RPCs (chPCs) is not maintained as a cycling population after OTX2 mutagenesis. Two of the most upregulated genes in the new cluster were BRINP2 and CDKN1A, both associated with cell cycle exit (Bunz et al., 1998; Kawano et al., 2004). This suggests that one of the functions of OTX2 in restricted RPCs is to promote the continuation of cell division for another one or two cell cycles. As previously described, THRBCRM1 restricted RPCs have gene expression profiles distinct from multipotent RPCs, including down-regulation of genes such as VSX2, whose loss-of-function phenotype suggests that it normally promotes the cell cycle (Burmeister et al., 1996; Buenaventura et al., 2018). Thus, in the transition from multipotent RPCs to restricted RPCs, part of the restricted RPC gene regulatory network may be responsible for promoting limited cell division. A proliferative role for OTX2 may also be reflected in loss of PH3 and EdU labeling in the RPE that we observed in OTX2^CRISPR^ mutant patches of the RPE, when OTX2 mutation was induced at E1.5.

While there is some signature of PR genes (subcluster 10-2), there are intriguing mixtures of gene expression profiles. Though this population seems definitively to not be dividing, there are signatures of genes normally enriched in dividing restricted RPCs. These include TFAP2D and OLIG2, which are robust markers of restricted RPCs in this dataset and from previous observations, respectively. Expression of a number of RGC-associated genes is upregulated in these cells, including the expression of TFAP2D, previously found to be expressed in RGCs (Li et al., 2014). One possible scenario is that these would-be PRs are trans-differentiating into RGCs and this alternative path to becoming an RGC is distinct from the normal path. In addition, fate trajectories obtained using each cell’s expression overlapping profiles also suggests that the majority of the cells in cluster 10 are following a predominantly RGC fate.

Taken together, we propose a model by which OTX2 serves as a key positive regulator of photoreceptor genesis from restricted RPCs, while repressing specific subtypes of other retinal fates (Fig. 7A). In addition, OTX2 mutation affects the cell cycle in several contexts, suggesting an additional role for OTX2 in proliferation (Fig. 7A). At the gene regulatory network level, OTX2 represses key transcription factors involved in non-photoreceptor cell types (Fig. 7B). We anticipate that further combined use of the single cell sequencing/CRISPR gene editing approach with variation of time, targeted genes, and labeled cell populations will provide a powerful genetic strategy to examine developmental gene regulatory networks.

## Acknowledgements

Support was provided by NIH National Eye Institute grant R01EY024982 (to M.E.) and 3G12MD007603-30S2 (CCNY). The content is solely the responsibility of the authors and does not necessarily represent the official views of the National Eye Institute, the National Institute On Minority Health and Health Disparities or the National Institutes of Health. Jeffrey Walker and Jorge Morales provided excellent technical support with flow cytometry and confocal microcopy experiments. We thank the members of the Emerson lab for support throughout the project and Cosmin Tegla and Estie Schick for critically reading the manuscript.

## Methods

### Subject Details

All experimental procedures were carried out in accordance with the City College of New York, CUNY animal care protocols. Fertilized chick eggs were obtained from Charles River, stored in a 16°C room for 0-10 days and incubated in a 38°C humidified incubator. Experiments were performed on unbiased number of female and male animals. For the scRNA sequencing experiment only, retinas from embryos PCR-identified as female were processed, to avoid variability from the reference genome.

For sex determination, genotyping PCRs were completed as explained in Clinton et al., 2001. Briefly, the technique is based on the presence of *Xho* I repeats, a class present on the W chromosome. A PCR reaction using primers flanking that region (forward 5’.CCCAAATATAACACGCTTCACT 3’; reverse 5’ GAAATGAATTATTTTCTGGCGAC 3’) as well as primers flanking a control region (forward 5’ AGCTCTTTCTCGATTCCGTG 3’, reverse 5’ GGGTAGACACAAGCTGAGCC 3’) were used. The presence of both bands identifies DNA of female origin, while presence of only the control gene band marks DNA of male origin.

Experiments were done in non-randomized, non-blinded conditions.

### CRISPR/Cas9-induced mutation design

To determine guide RNA sequences optimal for the ablation of the OTX2 gene www.chopchop.cbu.uib.no online tool was used, and the sequence of the first two coding exons of the gene were analyzed (Montague et al., 2014; Labun et al., 2016). The target DNA sequence with the best score calculated according to the online algorithm were selected and processed further.

The sequence of each guide RNA, excluding the PAM motif, was cloned into a modified px330 vector (Addgene number #42230) between the two *Bbs*I sites downstream the U6 promoter, as described in Cong et al., 2013. In the px330 vector, the CBh promoter was replaced with the CAG synthetic promoter (Okabe et al., 1997), using *Xba*I *Age*I restriction sites upstream of the humanized *S. Pyogenes* Cas9, herein called p18. Following plasmid delivery into target cells, Cas9 protein induces a double-stranded break which is resolved through the error-prone DNA repair mechanism, therefore genomic modifications were obtained through the non-homologous end joining pathway (NHEJ).

Expression vectors containing either florescent reporters or nuclear LacZ driven by the CAG promoter were used as electroporation controls (Matsuda and Cepko 2004; Buenaventura et al., 2018; Emerson et al., 2011). For the CAG::tdT reporter, the CAG promoter was cloned with *Sal*I *EcoR*I in the STATIA vector. This represents a modified version of the STAGIA3 plasmid, where EGFP was replaced with TdTomato using *Age*I *Bsr*GI restriction sites.

### Electroporation and explant culture

*In vivo* electroporation was performed at two developmental time points E1.5 (HH stage 9-11) and E3 (HH 18) on healthy-developed embryos. At E1.5, plasmids and Fast Green tracer (0.1% final concentration) in TE buffer was injected into the optic vesicle using a pulled glass needle. A sharp negative electrode was inserted into the forebrain at equal distance between the optic vesicles, while a mobile, positive electrode, was placed near the exterior side of the optic vesicle.

For E3 electroporations, DNA-dye mixes were injected into the subretinal space. The negative electrode penetrated the head in proximity to the eye’s dorsal side and the positive electrode was placed frontal to the eye. For both types of *in vivo* electroporations DNA cocktails contained the reporter plasmid used as electroporation control along with the p18 vector, with or without guide RNA sequence, all at a concentration of 2 µg/µl. Three pulses of 10V were applied using a Nepagene Super Electroporator NEPA21 Type II electroporator. Following the procedure, the windowed eggs were covered with transparent tape and returned to the 38°C incubator.

*Ex vivo* electroporations were done as previously reported in Buenaventura el al., 2018. Briefly, prior to electroporation the retinas were dissected into warm DMEM/F12 media (Life Technologies, 11320082). Plasmids of interest diluted in 1X PBS for a total volume of 50 µl filled the electroporation cuvette where the retina was placed, with the lens surface attached to the negative electrode. Five 50 ms pulses of 25V with a 950 ms interpulse interval were applied using the same electroporator. In general, plasmids were used at following concentrations – 100 ng/µl for the reporter plasmids driven by the CAG promoter, 160 ng/µl for enhancer-driven reporter plasmids and 200 ng/µl for Cas9 based plasmids. After electroporation, lenses were dissected out and retinas were placed on porous filters of 0.2 µm (13 mm Nuclepore Track-Etch Membrane Whatman filters) floating on 1ml basic culture media, 10% FCS, 1X Pen/Strep, 1X L-glutamine in DMEM/F12 (Life Technologies, 10378016) and incubated at 37°C with 5% CO_2_.

### Immunohistochemistry

Retinas processed for immunohistochemistry were fixed in 4% paraformaldehyde for 30 minutes at room temperature, sunk in 30% sucrose and snap-frozen in OCT (Sakura Tissue-Tek, 4583). Vertical sections of 20 µm were obtained using a Leica Cryostat and collected on microscopy slides (FisherBrand, 12-550-15).

Immunofluorescence protocol was as described in Buenaventura el al., 2018. A list of all primary antibodies used, along with their vendor information and concentration used is shown in Table 1. Following incubation with the primary antibody, retinal sections were washed in 1X PBS, 0.1 % Tween for a total of 30 minutes at room temperature, then blocked for additional 30 minutes in the blocking solution described above. Secondary antibodies used were: Alexa 488 and Alexa 647 to a 1:400 dilution (Jackson Immunoresearch, 115-605-207), and Cy3 to a 1:250 dilution (Jackson Immunoresearch 115-165-205). All secondary antibodies were used in different combinations according to the host of the primaries (anti-rabbit, anti-mouse, anti-sheep and anti-goat) and were appropriate for multi-labeling. For nuclear counterstaining a solution of 4′,6-diamidino-2-phenylindole (DAPI) in 1X PBS was applied on sections prior to three final washes of 15 minutes at room temperature in 1X PBS. Slides were mounted in Fluoromount (Southern Biotech, 0100-01) with 34×60mm cover slips (VWR, 48393 106).

To immunofluorescently label dissociated cells in suspension, the following procedure was applied – after the 15 minutes fixation retinas were washed with 1X PBS, followed by washes in 1X PBS, 0.1% Tween. Cells were blocked for 1 hour in 5% normal serum of the species the secondary antibodies were raised in, then incubated overnight at 4°C in a solution containing the primary antibodies and 5% serum. The next day, cells were washed with 3ml 1X PBS, 0.1% Tween and blocked in 5% serum for 30 minutes. Secondary antibodies were added to the blocking solution to the concentrations noted above and incubated for 1 hour at room temperature protected by light.

For EdU incorporation in ovo, a solution of 10 µM in 1X PBS was added on top of the embryo 5 hours prior to sacrifice (Warren et al., 2009). To develop, a Click-iT EdU Alexa Fluor 647 imaging kit was used in addition to the regular immunostaining protocol (Invitrogen, C10340).

### Microscopy

High magnification images were acquired with an Axiozoom V16 microscope using a PlanNeoFluor Z 1x objective at a digital zoom of 37.5.

Confocal micrographs were acquired at a 1024 x 1024 resolution using an inverted Zeiss LSM710 confocal microscope with an EC Plan-Neofluar 40x/1.30 Oil DIC M27 objective. During acquisition a ZEN software was used. All images were processed and converted into tiff. format using the FIJI version of ImageJ (Schneider et al., 2012). Figures were assembled using Affinity Designer vector editor; general brightness and contrast adjustments were done when necessary on the entire field imaged.

### Dissociation and cell sorting

Retinal explants were dissociated after being cultured for 48 hours with a papain-based protocol (Worthington, L5003126) as described in Jean-Charles et al. 2018. After dissociation, cells were fixed in 4% PFA at room temperature for 15 minutes, washed and analyzed, or immunostained as described above, prior to analysis.

The florescent activated cell sorting (FACS) of the GFP reporter was done using a BD FACS Aria machine with a 488nm laser. For sorting experiments requiring high cell viability post sorting and nucleic acid extractions, cells were resuspended and sorted in cold DMEM 10% FBS (Gibco 10437-010) media.

### Library preparation, sequencing and data analysis

Single cell RNA sequencing of OTX2ECR2::GFP positive cells in the OTX2^CRISPR^ mutants and WT conditions, 40,000 GFP positive cells from four retinas per sample were collected into DMEM 10% FBS media, pelleted at a speed of 500 *xg*, then resuspended in the same media. A trypan blue exclusion viability test confirmed that the percentage of viable cells was greater than 70%. Approximatively 12,000 cells were input into the 10X Genomics Chromium machine and individual droplets were formed containing cell and molecule specific barcodes. Libraries were sequenced on an Illumina 4000 sequencer. The data alignment on the GRCg6a reference genome, barcode processing and generation of cell-gene matrices was done at the Single Cell Analysis core facility at Columbia Genome Center using the 10X Genomics pipeline. Matrices were further analyzed with R using the Seurat package (Satija et al., 2015). For the analysis, the dataset obtained from the OTX2 WT retinas was processed individually, as well as in combination with the OTX2^CRISPR^ dataset, projects called “wt” and “merged”, respectively. Briefly, for both “wt” and “merged” projects, from the mitochondrial and ribosomal genes, genes detected and unique molecular identifiers (UMIs) outliers were filtered out from the downstream analysis. Cells were assigned cell cycle phases based on their gene expression, and the cell cycle genes were also regressed out. Genes used for the mitochondrial, ribosomal and cell cycle regressions are shown in Supplemental Table 2.

Data was normalized and scaled, and principal component analysis was performed based on a list of variable genes previously computed.

### Quantification and Statistical Analysis

Quantitative analysis of the florescent reporters and of the antibody-detected markers was done with a BD LSRII machine, using the 488nm, 561nm and 633nm lasers. The analysis was carried out with the FlowJo Version 10.2 software.

In each experiment for quantitative analysis, a minimum of four biological replicates were used in at least three technical replicates. Comparisons between OTX2^CRISPR^ mutants and control groups were done using student’s *t*-test with independent samples, after the Shapiro-Wilk normality test and Levene’s equality of variances assumptions confirmed the normal distribution of the data. For each quantification, one eye was used as experimental (OTX2^CRISPR^), while the contralateral represented the WT control. Error bars represent 95% confidence intervals (c.i.). For all experiments statistical analysis was done using JASP 0.9.0.1, GraphPad – PRISM and Microsoft Office Excel 16 software.

For the transcriptome analysis, four retinas were pooled for each of the WT and OTX2^CRISPR^ samples; for each of the mutant retinas included in the sample, the contralateral eye was used as WT control to limit the variability across individual animals.

