## Supplemental Information for "Single cell profiling of CRISPR/Cas9-induced OTX2 deficient retinas reveals fate switch from restricted progenitors"

### Supplemental Figure 1

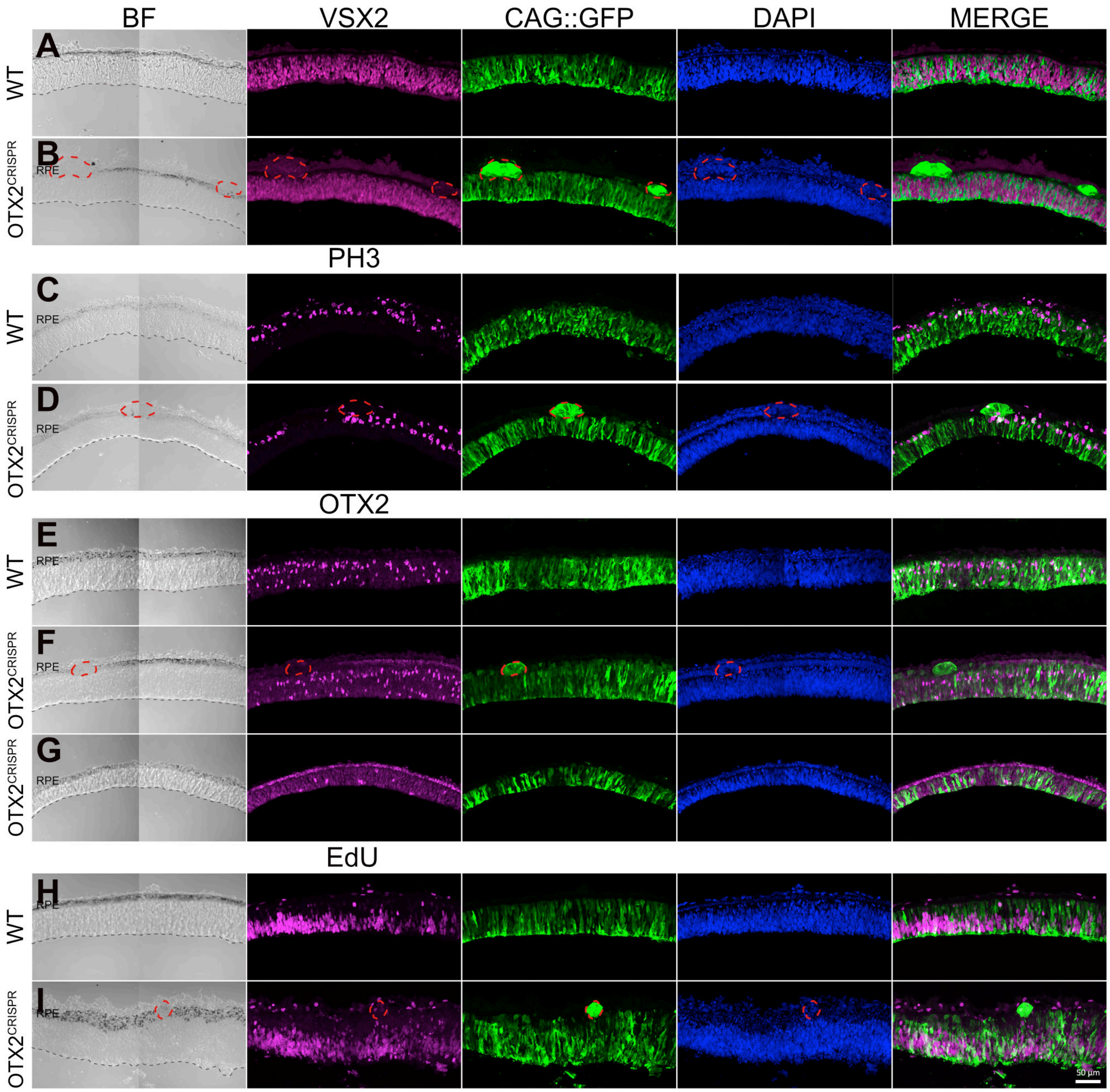

### Supplemental Figure 2

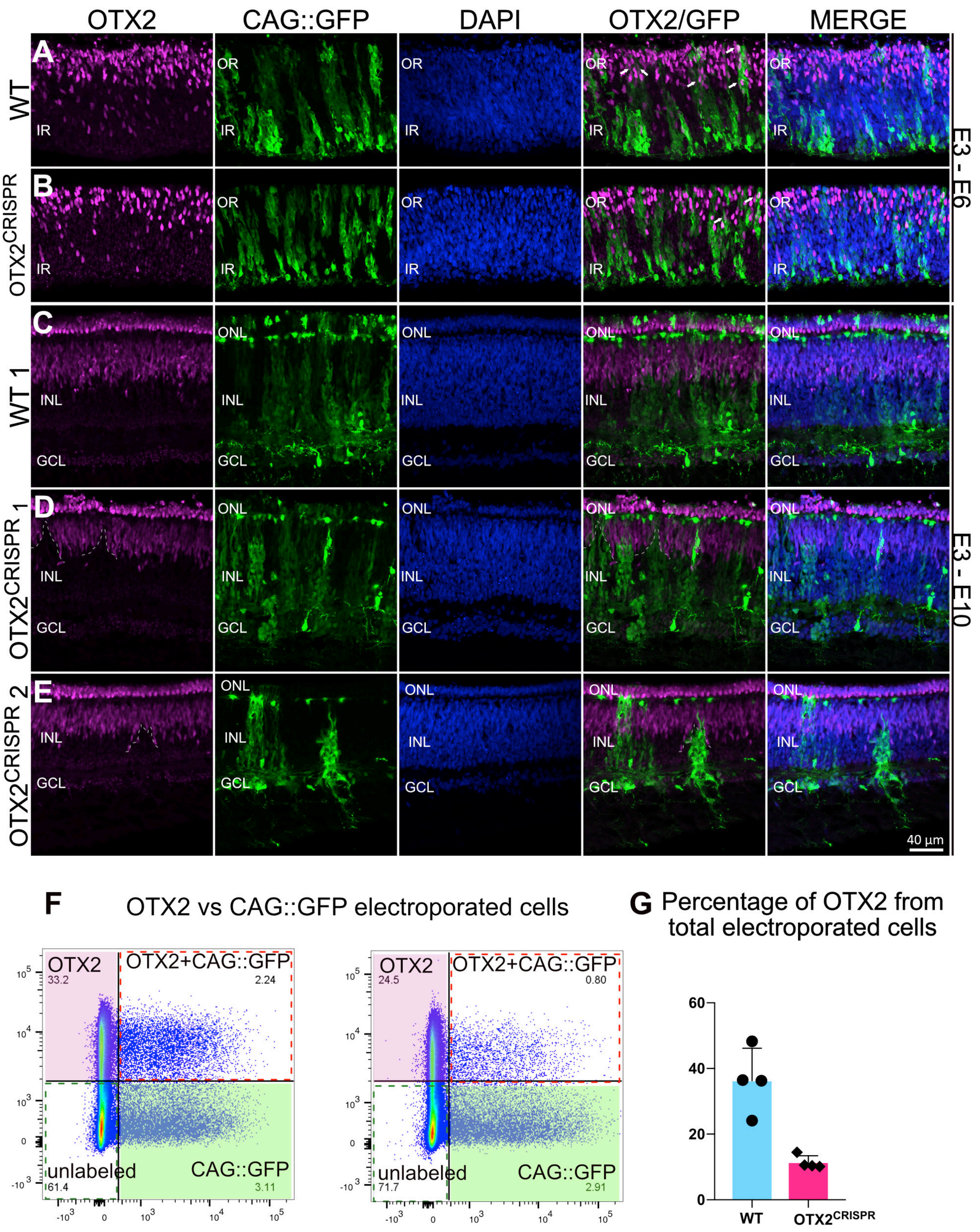

### Supplemental Figure 3

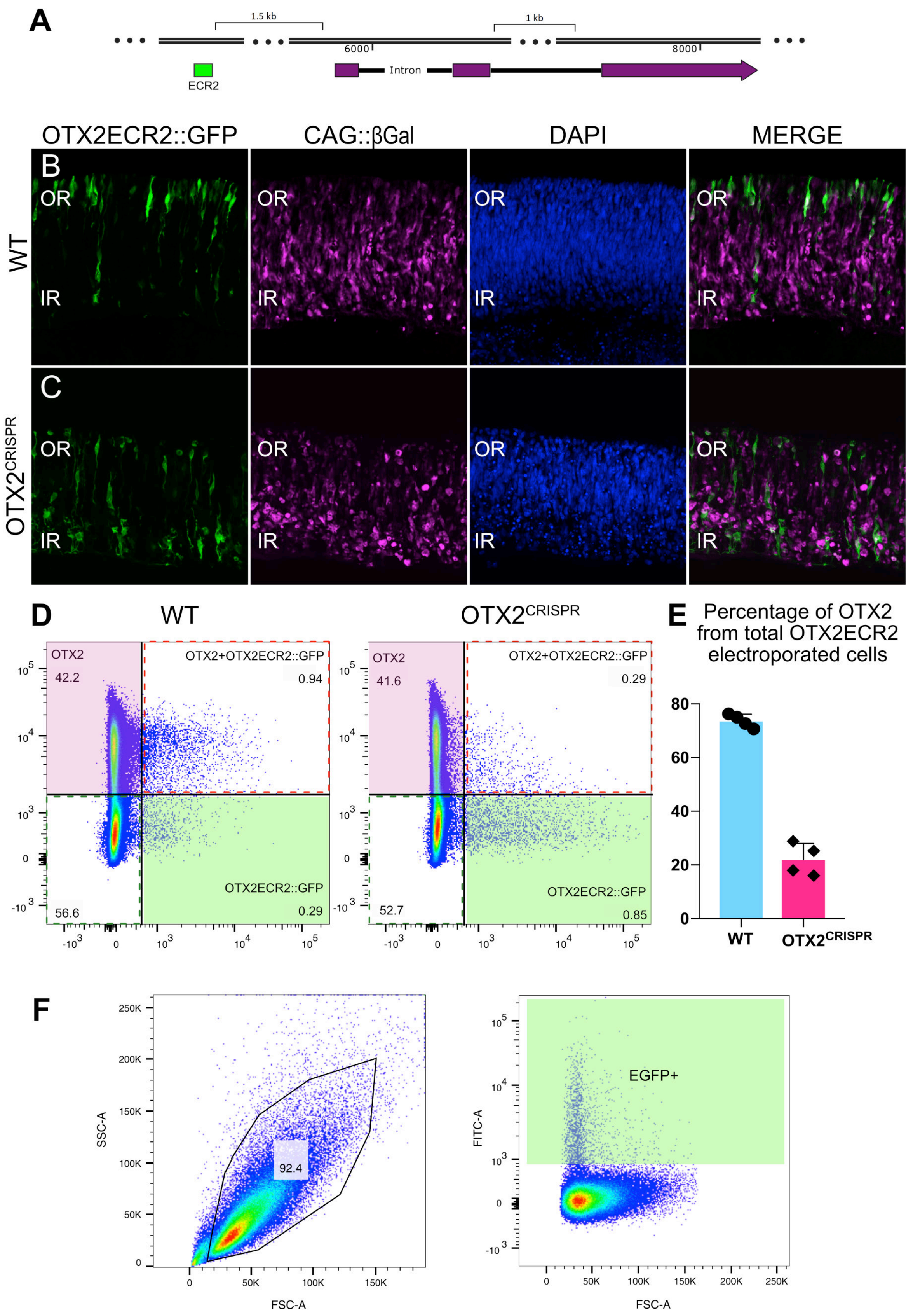

### Supplemental Figure 4

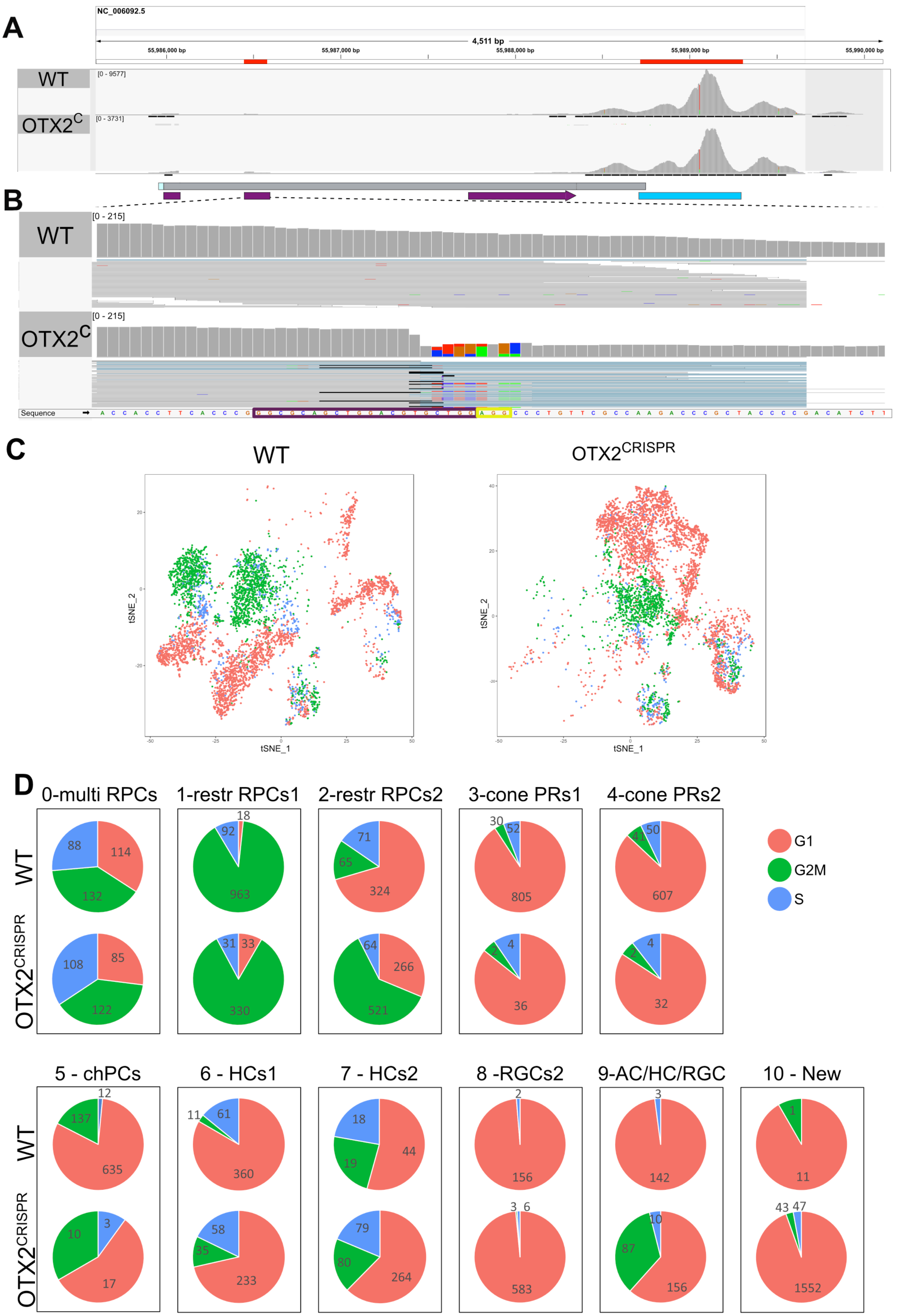

### Supplemental Figure 5

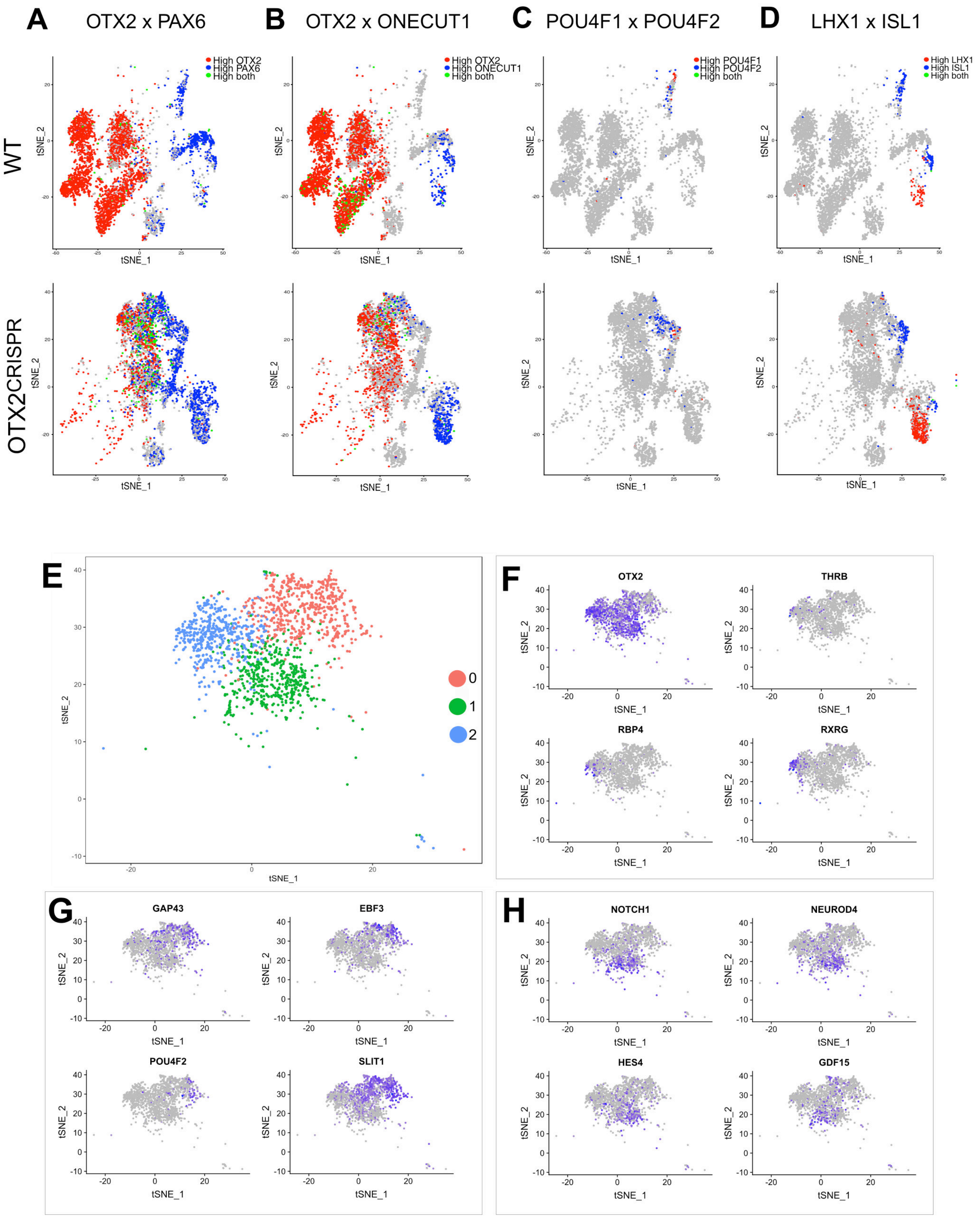

### Supplemental Figure 6

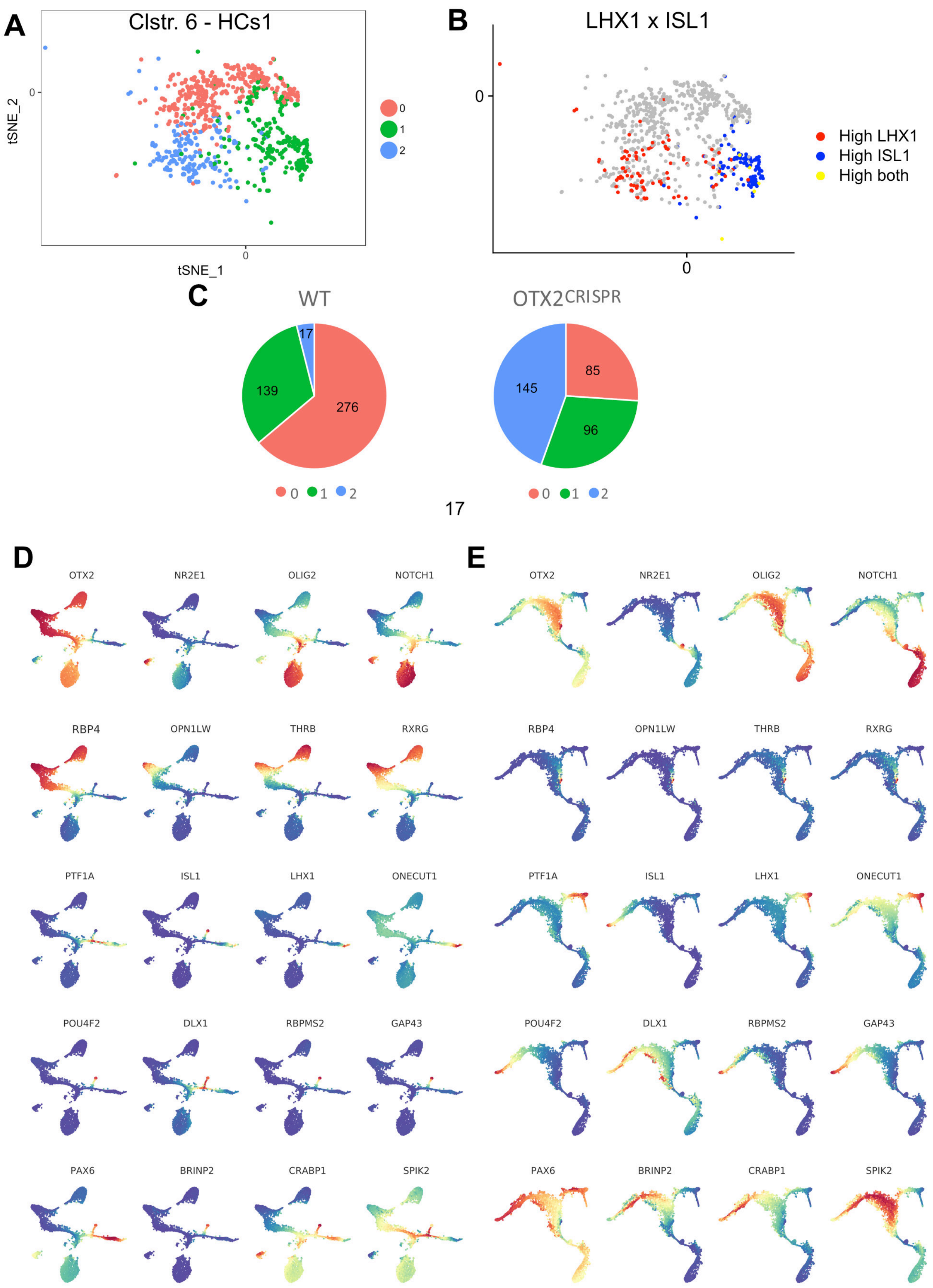

### Supplemental Figure 7

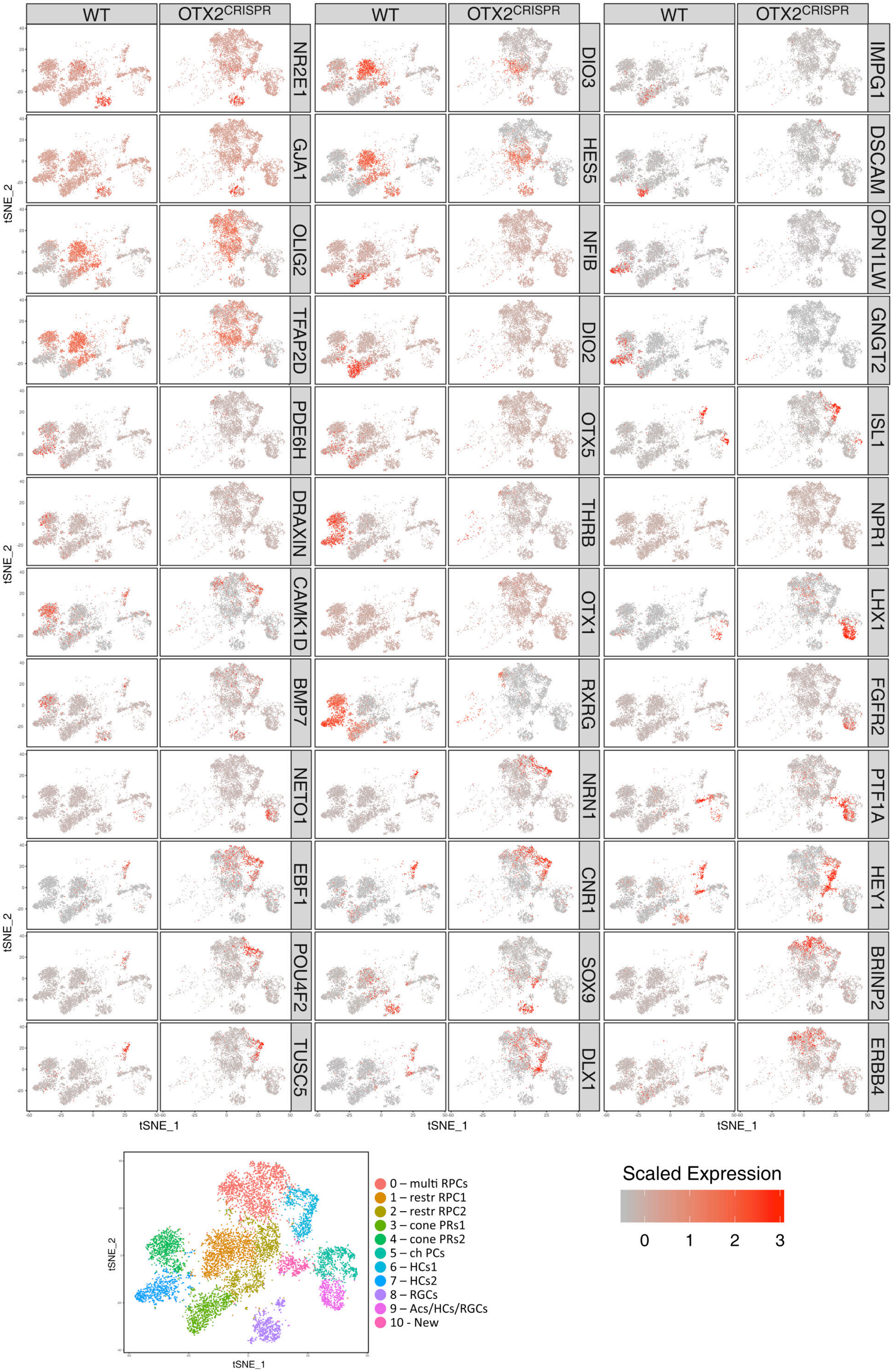

### Supplemental Table 1

| **Quantification** | **Figure** | **Average WT (±95% C.I.) (%)** | **Average $OTX2^{CRISPR}$ (±95% C.I.)** | **St. dev. WT** | **St. dev. $OTX2^{CRISPR}$** | **Number of biological replicates (n)** |
|---|---|---|---|---|---|---|
| OTX2 protein (Goat antibody)[1] | Figure 2D | 40.42±3.89 | 19.78±3.15 | 3.97 | 3.22 | 4 |
| PAX6 protein[1] | Figure 2F | 37.79±6.74 | 69.17±6.44 | 7.68 | 6.44 | 5 |
| THRBCRM2::GFP[1] | Figure 2H | 6.35±2.3 | 0.72±0.26 | 2.62 | 0.3 | 5 |
| OTX2 protein (Rabbit antibody)[1] | Suppl. Figure 2G | 36.29±967 | 11.39±2.01 | 9.87 | 2.05 | 4 |
| OTX2 protein (Goat antibody)[2] | Suppl. Figure 3E | 73.67±2.44 | 22.02±5.91 | 2.49 | 6.03 | 4 |

1. Percentages from total number of electroporated cells
2. Percentages from total number of OTX2ECR2::GFP electroporated cells

### Supplemental Table 2

**Mito**

| | |
|---|---|
| 1 | MTCH2 |
| 2 | MTCH1 |
| 3 | MTFR1 |
| 4 | MTHFD2 |
| 5 | MTHFD2 |
| 6 | MTG1 |
| 7 | MTERF3 |
| 8 | MTRF1 |
| 9 | MTHFD1L |
| 10 | MTFMT |
| 11 | MTO1 |
| 12 | MTPAP |
| 13 | MTG2 |
| 14 | MTFR2 |
| 15 | MTIF3 |
| 16 | MTIF2 |
| 17 | MTRF1L |

**Ribo**

| | | | | | | | |
|---|---|---|---|---|---|---|---|
| 1 | RPL18A | 21 | RPS6KA6 | 41 | RPL32 | 61 | RPL19 |
| 2 | RPS16 | 22 | RPL39 | 42 | RPL26L1 | 62 | RPL27 |
| 3 | RPS19BP1 | 23 | RPS3A | 43 | RPS14 | 63 | RPL36 |
| 4 | RPL3 | 24 | RPL34 | 44 | RPL3L | 64 | RPS28 |
| 5 | RPL24 | 25 | RPL7L1 | 45 | RPS2 | 65 | RPS15 |
| 6 | RPL8 | 26 | RPL9 | 46 | RPS15A | 66 | RPS26 |
| 7 | RPS6KA3 | 27 | RPS6KB2 | 47 | RPL6 | 67 | RPL17 |
| 8 | RPL31 | 28 | RPL27A | 48 | RPS5 | 68 | RPL37 |
| 9 | RPL21 | 29 | RPS13 | 49 | RPL12 | 69 | RPS6 |
| 10 | RPS3 | 30 | RPS6KL1 | 50 | RPL7A | 70 | RPS23 |
| 11 | RPS11 | 31 | RPS6KA5 | 51 | RPL35 | 71 | RPS19 |
| 12 | RPL15 | 32 | RPS29 | 52 | RPL38 | 72 | RPS7 |
| 13 | RPL14 | 33 | RPS24 | 53 | RPL23A | 73 | RPS4X |
| 14 | RPS20 | 34 | RPL37A | 54 | RPS6KB1 | 74 | RPL36A |
| 15 | RPL7 | 35 | RPL5 | 55 | RPS21 | 75 | RPL4 |
| 16 | RPL30 | 36 | RPS8 | 56 | RPL22 | 76 | RPL13 |
| 17 | RPS27A | 37 | RPL35A | 57 | RPS6KA1L | 77 | RPL29 |
| 18 | RPS6KC1 | 38 | RPL22L1 | 58 | RPL11 | 78 | RPL10A |
| 19 | RPS6KA2 | 39 | RPS17 | 59 | RPS25 | 79 | RPS10 |
| 20 | RPS12 | 40 | RPS27L | 60 | RPS27 | 80 | RPL23 |

**Cell cycle**

| | | | | | | | | | |
|---|---|---|---|---|---|---|---|---|---|
| 1 | MCM5 | 21 | GMNN | 41 | CHAF1B | 61 | CCNB2 | 81 | KIF23 |
| 2 | PCNA | 22 | WDR76 | 42 | BRIP1 | 62 | CKAP2L | 82 | HMMR |
| 3 | TYMS | 23 | SLBP | 43 | E2F8 | 63 | CKAP2 | 83 | AURKA |
| 4 | FEN1 | 24 | CCNE2 | 44 | HMGB2 | 64 | BUB1 | 84 | PSRC1 |
| 5 | MCM2 | 25 | UBR7 | 45 | CDK1 | 65 | KIF11 | 85 | ANLN |
| 6 | MCM4 | 26 | POLD3 | 46 | NUSAP1 | 66 | ANP32E | 86 | LBR |
| 7 | RRM1 | 27 | MSH2 | 47 | UBE2C | 67 | TUBB4B | 87 | CKAP5 |
| 8 | UNG | 28 | ATAD2 | 48 | BIRC5 | 68 | GTSE1 | 88 | CENPE |
| 9 | GINS2 | 29 | RAD51 | 49 | TPX2 | 69 | KIF20B | 89 | CTCF |
| 10 | MCM6 | 30 | RRM2 | 50 | TOP2A | 70 | HJURP | 90 | NEK2 |
| 11 | CDCA7 | 31 | CDC45 | 51 | NDC80 | 71 | CDCA3 | 91 | G2E3 |
| 12 | DTL | 32 | CDC6 | 52 | CKS2 | 72 | JPT1 | 92 | GAS2L3 |
| 13 | PRIM1 | 33 | EXO1 | 53 | NUF2 | 73 | CDC20 | 93 | CBX5 |
| 14 | UHRF1 | 34 | TIPIN | 54 | CKS1B | 74 | TTK | 94 | CENPA |
| 15 | CENPU | 35 | DSCC1 | 55 | MKI67 | 75 | KIF2C | | |
| 16 | HELLS | 36 | BLM | 56 | TMPO | 76 | RANGAP1 | | |
| 17 | RFC2 | 37 | CASP8AP2 | 57 | CENPF | 77 | NCAPD2 | | |
| 18 | RPA2 | 38 | USP1 | 58 | TACC3 | 78 | DLGAP5 | | |
| 19 | NASP | 39 | CLSPN | 59 | FAM64A | 79 | CDCA2 | | |
| 20 | RAD51AP1 | 40 | POLA1 | 60 | SMC4 | 80 | ECT2 | | |

**Supplemental Figure 1**. E1.5-induced OTX2 mutation triggers defects in both the RPE and retina.
Mutant retinas vary in thickness compared to the WT. VSX2 , PH3, OTX2 expression in WT (**A, C, E**) and OTX2$^{CRISPR}$ mutants (**B, D, F, G**).
**H-I** EdU S-phase labeling in the WT and OTX2$^{CRISPR}$ retinas. BF, brightfield; RPE, retinal pigment epithelium; OR, outer retina; IR, inner retina.

**Supplemental Figure 2**. Qualitative analysis of E3 *in vivo* electroporated retinas analyzed at E6 (**A-B**) and E10 (**C-E**).

**A-B**. White arrows denote electroporated cells that are positive for OTX2. **C-E**. GFP-positive, OTX2-negative patches (dotted lines) are present in the INL and PR layers of OTX2$^{CRISPR}$ mutants **(E)**. ONL, outer nuclear layer; INL, inner nuclear layer; GCL, ganglion cell layer.
**F**. Representative dot plots showing the overlap between OTX2 protein (y-axis) and CAG::GFP (x-axis) in WT and OTX2$^{CRISPR}$ dissociated retinas.
**G**. Percentage of OTX2-positive cells present within electroporated populations in WT and OTX2$^{CRISPR}$ retinas. Error bars represent 95% confidence intervals, p=0.0021, n=4 each WT and OTX2$^{CRISPR}$.

**Supplemental Figure 3**. OTX2ECR2::GFP reporter activity in WT and OTX2$^{CRISPR}$ retinas. Retinas were electroporated ex vivo at E5 with OTX2ECR2::GFP with Control or OTX2$^{CRISPR}$ plasmids and analyzed post 48 hours.
**A**. Schematic representation of the OTX2 genomic locus, and the location of the ECR2 element, 1.5kb upstream of the start codon.
**B-C**. OTX2ECR2::GFP reporter predominantly labels cells in the outer WT retina. This population is severely reduced in OTX2$^{CRISPR}$ mutants, and a new population located in the inner retina is formed. OR, outer retina; IR, inner retina.
**D-E**. Representative dot plots showing the overlap between OTX2ECR2::GFP and OTX2 protein. The population of OTX2ECR2::GFP-only cells is increased in OTX2$^{CRISPR}$ retinas. Error bars represent 95% confident intervals. p<0.001, n=4.
**F**. FACS plots showing the collection gates for the cells sorted prior to single cell library preparation.

**Supplemental Figure 4**. Analysis of the OTX2 reads aligned to the *Gallus gallus* genome, GRC6 assembly (GCF_000002315.5) using IGV (Integrative Genomics Viewer) **(A-B)** and distribution of cell cycle phases across the two samples **(C-D)**.
**A**. View of the OTX2 locus. First red box represents the second coding exon, while the last one represents the 3’ end of the gene. The majority of the reads are located at the 3’ end, due to the specifics of the technique.

**B**. Increased magnification of the second coding exon. Grey vertical bars represent the reads aligned to that region of the gene. Purple box around the sequence highlights the Cas9 target DNA and the yellow box highlights the PAM motif. Colored vertical blocks in the $OTX2^{CRISPR}$ alignment represent the mismatches identified in the aligned reads; black horizontal fine lines show deletions of the aligned reads.
**C**. TSNE plots of the 11 clusters of the WT (left) and $OTX2^{CRISPR}$ (right) cells.
**D**. Percentages of cell cycle signatures per clusters across the two datasets.

**Supplemental Figure 5**. Overlap between different markers (**A-D**) and characterization of cluster 10-New.

TSNE plots showing the overlap between PAX6 (**A**) or ONECUT1 (**B**) in WT and $OTX2^{CRISPR}$ cells. **C-D**. TSNE plots showing the overlap between POU4F1 and POU4F3 (**C**) and ISL1 and LHX1 (**D**) in WT and $OTX2^{CRISPR}$ cells.

**E**. Subclustering of cluster 10 using 9 PCs revealed 3 distinct subclusters (0-2). **F-H**. OTX2 expression in cluster 10 and markers of cluster 2 (**F**), cluster 0 (**G**) and cluster 1 (**H**).

**Supplemental Figure 6**. Characterization of cluster 6-HCs1 and markers for the assignment of pseudotime states.
**A-B**. Subclustering cluster 6-HC1 leads to 3 subclusters (0-2). B. Distribution of LHX1 and ISL1 within cluster 6.
**C**. Pie chart showing percentage of cells in cluster 6 subclusters in WT and $OTX2^{CRISPR}$ samples. **D-E**. Expression of different markers in the pseudotime model for WT (**D**) and $OTX2^{CRISPR}$ cells (**E**).

**Supplemental Figure 7.** Representative markers for each cluster in WT and $OTX2^{CRISPR}$ samples.

**Supplemental Table 1**. Quantitative analysis of the $OTX2^{CRISPR}$ mutants with different markers using flow cytometry.

**Supplemental Table 2**. Genes used for the mitochondrial (Mito) and ribosomal (Ribo) regression, and for cell cycle scoring.
